# Multiplexed experimental strategies for fragment library screening using SPR biosensors

**DOI:** 10.1101/2020.12.23.424167

**Authors:** Edward A. FitzGerald, Darius Vagrys, Giulia Opassi, Hanna F. Klein, David J. Hamilton, Pierre Boronat, Daniela Cederfelt, Vladimir O. Talibov, Mia Abramsson, Anna Moberg, Maria T. Lindgren, Claes Holmgren, Doreen Dobritzsch, Ben Davis, Peter O’Brien, Maikel Wijtmans, Jacqueline E. van Muijlwijk-Koezen, Roderick E. Hubbard, Iwan J.P de Esch, U. Helena Danielson

## Abstract

Surface plasmon resonance biosensor technology (SPR) is ideally suited for fragment-based lead discovery. However, generally suitable experimental procedures or detailed protocols are lacking, especially for structurally or physico-chemically challenging targets or when tool compounds are lacking. Success depends on accounting for the features of both the target and the chemical library, purposely designing screening experiments for identification and validation of hits with desired specificity and mode-of-action, and availability of orthogonal methods capable of confirming fragment hits. By adopting a multiplexed strategy, the range of targets and libraries amenable to an SPR biosensor-based approach for identifying hits is considerably expanded. We here illustrate innovative strategies using five challenging targets and variants thereof. Two libraries of 90 and 1056 fragments were screened using two different flow-based SPR biosensor systems, allowing different experimental approaches. Practical considerations and procedures accounting for the characteristics of the proteins and libraries, and that increase robustness, sensitivity, throughput and versatility are highlighted.

## Introduction

The first methods and concepts of fragment-based drug discovery (FBDD) emerged over 20 years ago, and its subsequent adoption by practitioners continues to increase. Its core principles are accepted as viable means for finding and evolving hits for a chemical biology or drug discovery project^1–5^. FBDD, in its essence, is a reductionist alternative to high-throughput screening (HTS), built on the theory that a much broader chemical space can be more efficiently probed by using structurally diverse compounds with lower molecular weight than one would conventionally find in a high-throughput screening (HTS) or drug-like compound library. Fragment libraries therefore typically contain compounds with a molecular weight below *ca*. 300 Da and comprise hundreds to thousands, rather than hundreds of thousands, of compounds as for HTS. The small size of fragments and their chemical diversity makes it possible to identify novel scaffolds, alternative binding sites and modes-of-action in the early stages of a discovery program. But it has the disadvantage of providing only a few possible intermolecular contact points, exhibited as typically weak and transient interactions with their targets. They therefore have to be detected using very sensitive biophysical methods that monitor the interaction as such, although some indirect methods are also used. The methods must allow relatively high concentrations of fragments to be screened without running into experimental artifacts. In order of popularity,^6^ the methods currently used include X-ray crystallography^7^, Nuclear Magnetic Resonance (NMR), Surface Plasmon Resonance biosensor technology (SPR), thermal shift and *in silico* methods, functional screening and Isothermal Calorimetry (ITC).^8–10^ They differ in their sensitivity, capability of providing kinetic or structural information, experimental flexibility and risk of experimental artifacts. Their suitability for a certain target or project therefore varies. Since identified hits should be confirmed by an orthogonal method, at least two methods are required for hit discovery. Evolution of hits into leads can often be initiated by acquiring commercially available compounds (aka “analogue-by-catalogue”), but is later structurally guided using medicinal and computational chemistry approaches^8,9,11,12^.

FBDD has been a truly transformative approach, proven to be effective for the discovery of novel therapeutics and with various examples in the clinic, including four FDA-approved drugs to date: vemurafenib^13^, erdafitinib^14^, venetoclax^15^ and pexidartinib.^16^ Now that fragment libraries have become commercially available and screening technologies cheaper to acquire and implement, FBDD is a realistic option for smaller pharmaceutical companies and contract research organizations (CROs), as well as for academic core facilities and research groups. Nevertheless, success relies heavily on the capability to use sophisticated biophysical instrumentation and access to high quality protein samples and chemically diverse fragment libraries. The experimental strategy must be defined on a case-by-case basis, driven by a good understanding of the target characteristics and the types of lead compounds and modes-of-action that are of interest.

Time-resolved SPR biosensor-based interaction analysis was one of the earliest methods adopted for FBDD, with a sensitivity and throughput suitable for fragment library screening.^17–20^ Its versatility and information rich data output makes it useful also for supporting the evolution of hits and optimisation of leads, *i.e*. all the way to nomination of candidate drugs. Its use has increased over time and experimental strategies and applications are continuing to evolve. The method requires high quality target samples, but in lower quantities than for other methods. However, to fully exploit the possibilities, target variants and tool compounds should be available as controls to ensure that prepared sensor surfaces are functional and interact with ligands in an expected manner. Experiments need to be designed and evaluated using appropriate data analysis procedures, tailored specifically for each project.

New SPR-based biosensors developed specifically for FBDD, with increased sensitivity, throughput and features allowing experiments to be designed for fragment-based work have appeared on the market. However, detailed protocols or descriptions of generally suitable experimental procedures are lacking and disclosed methods for more challenging targets is limited.

Herein, we outline novel SPR-driven strategies for identifying fragment hits, the first critical step in a FBDD project. Two flow-based SPR biosensors with different features, two fragment libraries of different sizes and a panel of five target proteins (nine variants in total) representing structurally or physicochemically challenging targets and/or lacking tool compounds, was used to illustrate multiplexed experimental strategies (Fig. 1). The panel represents different target classes, varying in size and structural complexity, and availability of tool compounds. The results show that the range of targets amenable to an SPR biosensor-based approach is considerably expanded by adopting a multiplexed strategy, and hits can be identified even when using a relatively small library.

**Fig. 1.**
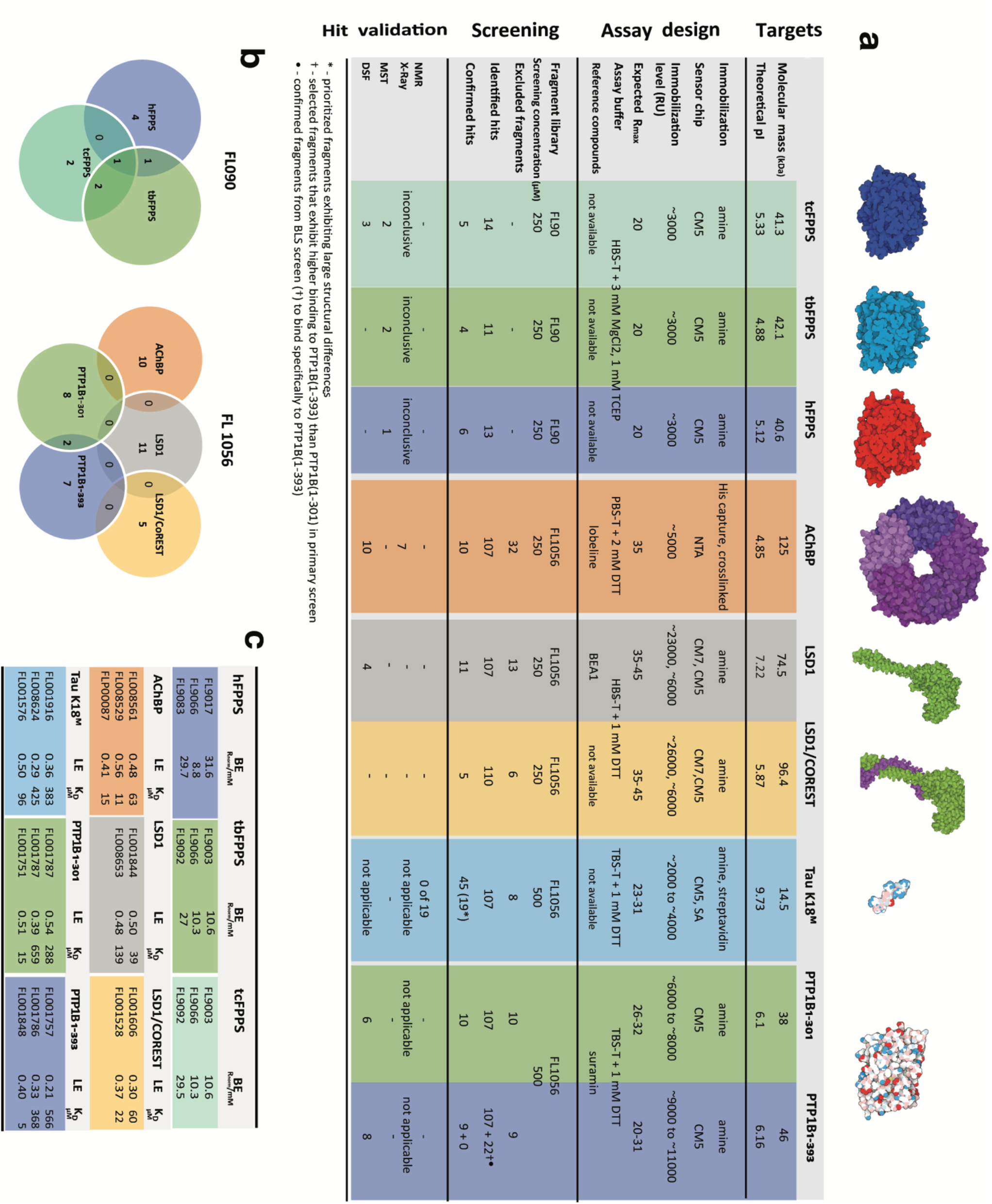
Overview of target proteins, fragment libraries and methods used for screening and validation of hits. **a,** Targets and their structural characteristics: Farnesyl pyrophosphate synthase (FPPS) from human, *trypanosoma brucei* and *cruzi* (h, tb and tc, respectively), acetylcholine binding protein (AChBP), lysine demethylase 1 (LSD1) with and without the protein cofactor COREST, tau K18^M^, catalytic domain of protein tyrosine phosphatase 1B (PTP1B) in two different lengths. Assay design with experimental details for SPR experiments. Screening strategy and results from different steps. Hit validation specifying orthogonal methods used and the current status. **b,** Venn Diagram highlighting identified hits for (left) FL90 and (right) FL1056. **c,** Table describing LE, BE and *K*D^app^ of confirmed fragment hits.

## SPR biosensor-based interaction analysis of fragments

When adopting time-resolved SPR biosensor systems for analysis of fragments, there are some unique challenges. Fragments are injected in a continuous flow of buffer in a microfluidic flow system containing a derivatized gold sensor surface to which the other molecule has been immobilised, embodying the sensor surface (Fig. 2a). By continuously monitoring the angle where the intensity of the reflected light has a minimum, due to surface plasmon resonance, it is possible to detect changes taking place at the surface in real time. Signals depend on the refractive index of the medium close to the gold layer and are dominated by interactions between analytes and immobilised molecules.

**Fig. 2.**
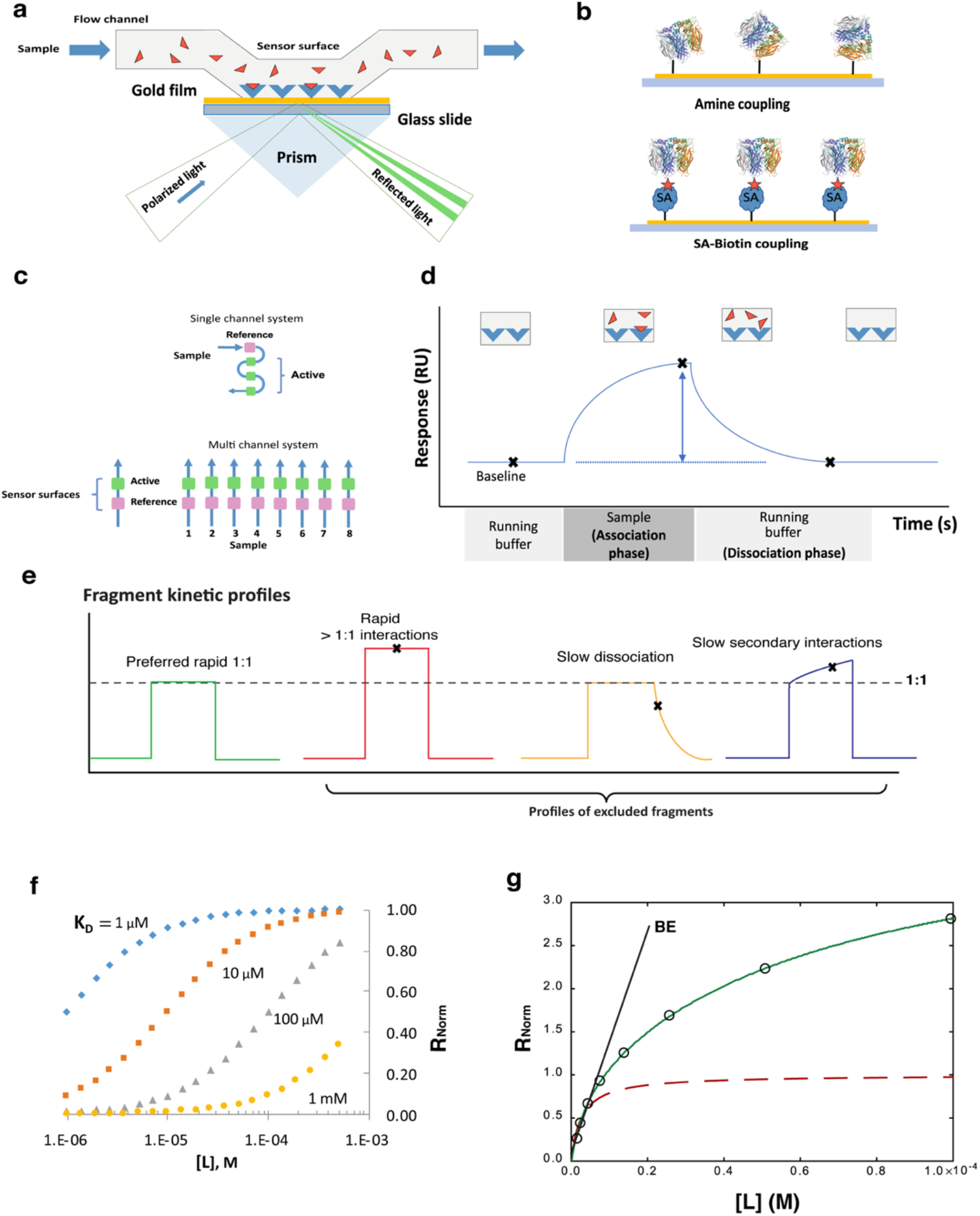
Overview of SPR biosensor technology principles and data. **a,** Graphical representation of SPR working principle. **b,** Visualization of targets immobilized to sensor surfaces in random orientation via covalent amine coupling (top) and in fixed orientation via non-covalent SA-biotin conjugation (bottom). **c**, Illustration of alternative layouts of flow systems and sensor surfaces. **d**, Depiction of events observed during an experiment. First, the system is equilibrated using running buffer and a baseline signal representing the surface with free target (P) is recorded. Binding events are observed as an increased signal upon the injection of a sample (L) over the derivatized surface during the association phase. The signal represents the ligand target complex (PL) whose concentration is defined by the concentration of injected analyte and the association rate constant *k*_on_ (units s^-1^). The dissociation of bound molecules is observed during the dissociation phase, when only running buffer flows over the surface. The curvature is defined by the dissociation rate constant *k*off (units M^-1^ s^-1^). Rate constants and the equilibrium dissociation constant *K*_D_ are quantified from sensorgrams recorded for a series of analyte concentration, using global non-linear regression analysis. *K*_D_ values can also be estimated from steady state data for concentration series. **e**, Binding behaviour markers: green – no artefactual binding behaviour, yellow – slow dissociation, blue – slow association, red – *R*>*R*_max_, −multiple artefactual binding behaviours. **f,** relationship between fractional occupancy at different screening concentrations and *K*_D_ values. **g,** Graphs representing *R*_norm_, *i.e*. the signal at steady state (*R*_eq_) normalized with respect to maximum binding level (*R*_max_). A simple reversible 1:1 interaction can be described by the Langmuir equation (red dotted line). For higher order interactions, representing interactions with multiple sites (2:1, 3:1 etc.) or non-specific interactions (≫1:1), signals higher than 1 will be seen (green line). The binding efficiency (BE) is estimated from the initial slope of the graph, representing the total binding to the target at low concentration.

### Sensor surface preparation

Due to the small size and rapid and weak affinities of fragments, successful SPR biosensor-based screening of fragment libraries requires stable and very sensitive sensor surfaces. In practice, assay development is a trial-and-error process where success relies also on practical considerations, such as the availability of high-quality target protein samples and the possibility to control the functionality of sensor surfaces. It is often necessary to produce target proteins specifically constructed for SPR biosensor analysis, increasing stability or introducing residues or tags for immobilisation.

Target proteins are immobilised to sensor chips pre-coated with a carboxylated polymer, varying in density, degree of carboxylation and chain length using well-established chemical and biological approaches, such as covalent amine or thiol coupling, capture via His-tags or biotin groups proteins using chelating groups (NTA), antibodies or streptavidin covalently attached to the surface. Coupling techniques that directly immobilize the target protein are preferred over the use of antibodies or other protein constructs as these additional proteins increase the density on the sensor surface and can lead to artifacts (*e.g*., interaction of fragments with non-target binding sites). When screening fragments against large proteins (>100 kDa), it can be advantageous to use polymer surfaces with a higher degree of carboxylation and a denser matrix. Targets can be immobilised in a random or defined orientation (Fig. 2b) and further modification of surfaces are possible using biological and chemical modification methods, either before or after immobilisation of target proteins. The orientation is not critical for FBDD, but potentially relevant binding sites should not be blocked. Multiple immobilisation strategies often have to be explored since it is critical to have a sensitive sensor surface with a fully functional target throughout the experiment.

### Microfluidic flow systems

State-of-the-art flow-based instruments vary in the number of channels in the microfluidic flow system and the number and relative position of sensor surfaces (Fig. 2c). This influences the flexibility of the experimental design and the throughput of assays. Instruments that allow the use of multiple sensor surfaces in a single flow channel are advantageous for advanced referencing of data, *e.g*. using multiple target variants or off-targets, while screening efficiency can be higher with systems that have multiple flow channels. For fragment analysis, it is necessary to use at least two surfaces in order to discriminate the typical weak binding to the properly folded and functional target from nonspecific binding to unfolded or otherwise non-functional forms of the target, immobilised contaminants or the matrix.

### Data output and assay sensitivity

The output from experiments is in the form of sensorgrams, representing response signals in resonance units (RU) and as a function of time (Fig. 2d). The detection limit depends on instrument features, how the assay is set up and the characteristics of the sensor surfaces used. The theoretical maximal response (*R*_max_) that can be expected for an interaction is dependent on the analyte binding capacity of the surface. By assuming that the target is fully functional and that the interaction follows a reversible 1-step mechanism with a 1:1 stoichiometry, it can be estimated as:

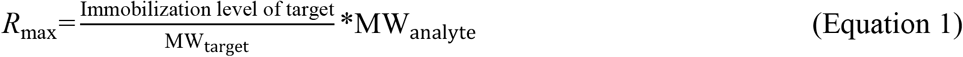

*R*_max_ can be determined experimentally by injecting a saturating concentration of a tool compound with a well-defined and reversible interaction with the target. Comparisons of theoretical and experimental *R*_max_ values define the degree of surface functionality and are important sensor surface quality controls, critical when establishing new surfaces and ensuring that surfaces do not deteriorate during a screen and can be used to predict if surfaces have adequate sensitivity for fragment studies.

To optimise the sensitivity of a surface, it is tempting to maximise the immobilization level of the target since it directly affects *R*_max_ (Equation 1). But this may result in mass transport problems and suboptimal surface characteristics. It is more important to use surfaces with pure and fully functional protein which also avoids the risk of false positives resulting from interactions with unfolded, denatured or aggregated protein. Selection of experimental conditions with respect to protein stability for the duration of experiments is particularly important for fragment screening where concentrations are high and specificity is low.

The signal expected for fragment screening hits is lower than *R*_max_ since they typically have low affinities and do not saturate the target, even when screening is performed at the highest practical concentration. Instead, for fragments with rapid interactions, the measured response corresponds to the equilibrium response (*R*_eq_) which depends on its affinity for the target (expressed as the equilibrium dissociation constant *K*_D_) and the concentration injected (L):

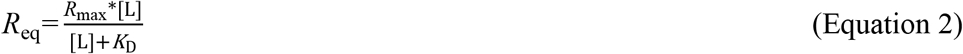

*K*_D_ can be defined either as the ratio of the concentrations of free target (P), free ligand (L) and the target-ligand complex (PL) at equilibrium, or as the ratio of the association and dissociation rate constants (*k*_on_ and *k*_off_, respectively):

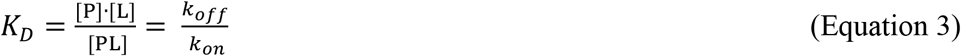

For successful fragment library screening, *R*_eq_ should be much higher than the detection limit for the system to be used. Due to the low molecular weight of fragments (affecting *R*_max_) and their weak and transient interactions with the target (affecting *K*_D_), signals are very low. The possibility to maximise *R*_eq_ by using high fragment concentrations ([L]) may in practice be limited by fragment availability, but other issues are common. Firstly, the relative contribution of secondary (non-specific) interaction with the target or sensor surface to the signal increases with concentration, making it essential to use as low concentrations of the fragments as possible. Secondly, inadequate fragment solubility in the assay buffer can result both in a lower effective concentration and aggregation on the surface thus potentially blocking the binding of subsequently injected fragments and obscuring the detection of hits. Increasing the solubility by increasing the solvent (DMSO) concentrations is not easily done since DMSO affects signals by changes to the refractive index as well as the functionality of targets. It is consequently essential to be accurate when pipetting samples in DMSO and to optimise and match running and sample buffers with respect to DMSO concentrations. Multiple controls and data correction routines are critical to detect and avoid solubility problems.

### Time-resolved analysis of fragments – kinetics

For many biomolecular interactions, sensorgrams have a clear curvature in both the association and dissociation phases (Fig. 2d). For fragments, the interactions are typically very rapid and sensorgrams are essentially square with no discernible curvature in the association or dissociation phases (Fig. 2e). However, data often show complexities due to rapid non-specific interactions with multiple sites (≫1:1 stoichiometry), slow dissociation or slow association to secondary sites. Depending on the target and goals of experiments, these non-ideal kinetic profiles can be more or less problematic. Square sensorgrams (Fig. 2e, left) are often considered as ideal for fragments, but only if the signal does not exceed theoretical *R*_max_ (red). Fragments showing a slow dissociation phase (yellow) are often excluded when selecting hits, but relevant fragments may thereby be overlooked. Similarly, slow association (blue) is often associated with problematic secondary/non-specific binding, although fragments interacting rapidly with single binding sites at low concentration and secondary sites at high concentrations will be excluded.

### Fragment library screening

Screening is efficiently done by injecting fragments at a single concentration and selecting hits based simply on the basis of signal levels at a defined time after injection, with controls taken before injection of samples and after a certain dissociation time (double base line controls). For SPR biosensor-driven screening it is essential that fragments have a high solubility under the experimental conditions and do not block the surface or result in carry over between injections. A pre-screening routine to identify compounds that may give rise to artifacts under the selected screening conditions and to confirm that the assay is sensitive and robust, is recommended. An option is to run the actual screen in forward and reverse order, and look for differences in signals, thus identifying problematic fragments and a deteriorating surface.^17^

To reliably identify fragments, screening should be done with sensitive assays that can discriminate low *R*_norm_ values from baseline and at concentrations higher than the expected *K*_D_ values, thus maximising *R*_norm_. Still, not much higher, due to the risk of signals arising from non-specific interactions and super stoichiometric binding. Fragments giving rise to square-shaped sensorgrams are prioritized, reflecting the typical fast on/fast off kinetics of fragment interactions with target proteins (Fig. 2e). However, fragments with deviations from rapid 1:1 interaction resulting in slow association, slow dissociation, or *R*_eq_>*R*_max_ (illustrated in Fig. 2e), or other secondary effects should not immediately be rejected as they may represent mechanistically interesting interactions. The number of expected hits depends on the size and quality and relevance of the library screened. It is often practical to select the 10% “best hits” based on signal levels and sensorgram shapes.

### Hit validation

Hits are validated in a follow-up experiment where hits are injected in a concentration series. Sensorgram shape and concentration dependence are evaluated. Signals for report points taken at steady state and different concentrations should ideally result in a hyperbolic saturation curve and be below *R*max, as described by Equation 2. To establish the interaction mechanism and estimate a *K*_D_-value, it is essential to use an analyte concentration range that is ≫ *K*_D_. For fragments, the affinity is often too weak for a practically useful concentration to be used (see Fig. 2f). Approximate *K*_D_ values (*K*_D_^app^) can be estimated by non-linear regression analysis using a 1:1 interaction model if the *K*_D_ is not much higher than the highest concentration used for screening (typically μM range), providing that *R*_max_ is not exceeded. When secondary/non-specific interactions occur, signals can be higher than *R*_max_ (Fig. 2g, solid curve). For very weak interactions, the relationship between signal and analyte concentration can be essentially linear, with only a slight curvature. It is then useful to estimate the Binding Efficiency (BE), which provides a measure of the ability of the fragment to bind to the target, without assuming an interaction model or stoichiometry.^21^ BE is determined from the initial slope of the signal *vs*. concentration graphs, *i.e*. at very low ligand concentrations where the amount of complex (*R*_eq_) is linearly dependent on BE, at different compound concentrations (*L*) (Fig. 2g, tangent).

## Results

### Selection and characterization of fragment libraries

Two fragment libraries with different features influencing the experimental strategies for SPR-based screening were used. Fragment library FL90 was repurposed from a commercially available library for screening by crystallography and was seen as a suitable alternative for practitioners who may not have access to an in-house library or specialized infrastructure for compound handling. It consists of 90 fragments characterized by a large chemical diversity and high solubility, which makes them ideally suited for crystallography.^22^ Fragment library FL1056 was comprised of 1056 fragments collated from SciLifeLab^23^ and FragNet compound collections. FragNet is a pan European initial training network in fragment-based drug discovery, with a special focus on the synthesis of 3D fragments, *i.e*. compounds that are not flat, but with a more complex structure. The FragNet fragments were selected on the basis of key physicochemical properties, including heavy atom count (HAC), molecular weight and lipophilicity (cLogP) were determined and compared to guidelines put forward by Astex for typical fragments (MW < 300 Da, ClogP < 3, HBA ≤ 3, HBD ≤ 3).^24^

The two libraries were profiled with respect to the 3D shape of fragments, assessed by principal moments of inertia (PMI)^25^ (Fig. 3). The analysis showed that FL90 (Fig. 3a) exhibited a significantly lower spatial complexity than FL1056 (Fig. 3b), consistent with the inclusion of specifically designed 3D fragments in FL1056. In addition, the expected Ligand Efficiency (LE) values based on potential *K*_D_ values were calculated for FL1056 (Fig. 3c). The table shows the LEs that can be expected for FL1056 library hits, depending on the HACs and *K*_D_ values. When screening the library at 250-500 μM, only fragments with *K*_D_ <1 mM can result in *R*_norm_-values > 50% (Fig. 2f).

**Fig. 3.**
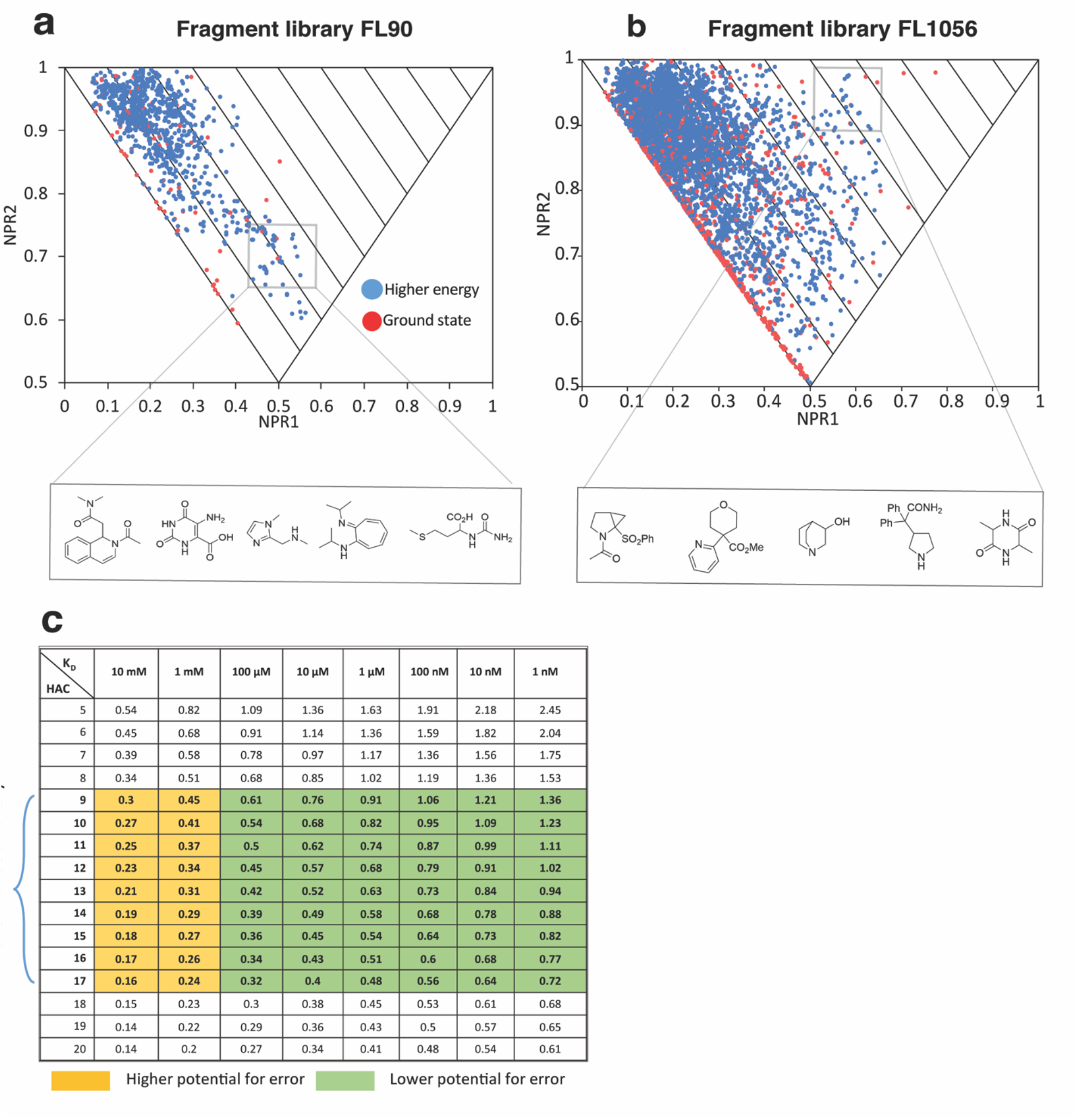
Analysis of structural diversity of fragment libraries and potential ligand efficiencies (LE) for fragments. Principal moments of inertia (PMI) analysis for all conformations up to 1.5 kcal mol-1 above the energy of the ground state conformation for each fragment. Triangular PMI plots show fragments with disc shapes at the bottom, rod shapes at the top-left and spherical shapes at the top-right. Conformations that lie furthest from the diagonal rod-disc axis have the most complex 3D shape as they deviate the most from planarity. **a**, FL90 (Frag Xtal Screen library, JENA Bioscience) and **b**, FL1056, selected from SciLifeLab^23^ and FragNet compound collections. Red dots indicate ground state conformers and blue dots show higher energy conformers. **c**, Expected Ligand Efficiency (LE) values calculated for FL1056, accounting for size (defined as Heavy Atom Count, HAC) and potential *K*_D_ values at 25 °C.

### Target production, assay development and quality controls

The procedures used to produce high quality protein for SPR experiments, as well as for handling and immobilisation of proteins, and validation of sensor surfaces, are detailed in methods and summarised in Fig. 1. Differential scanning fluorimetry (nanoDSF) was convenient for routine assessment of the structural integrity of new batches of protein and samples subjected to storage and handling and for identifying buffer conditions suitable for immobilisation and interaction experiments (Supplementary Table 2). However, it could not be used for disordered proteins, and tau K18^M^ was excluded also since lacks Trp and Tyr residues.

Assay development involved exploration of different coupling techniques and experimental conditions. Control experiments for evaluating sensor surfaces were performed for targets for which reference compounds available, *i.e*. AChBP, LSD1 and PTP1B variants (Supplementary Figs. 1 and 2). The sensor surfaces had required sensitivity for detecting interactions with low molecular weight compounds and stability to be used for 48 h, the maximum time used for a screening campaign.

### Experimental designs for single- and multi-channel systems

The two fragment libraries were screened using different biosensor systems and experimental approaches (Fig. 2c), with respect to the required experimental design and throughput. A single flow channel system was initially used for screening of FL90. It allowed screening simultaneous against three FPPS variants, thus allowing a direct specificity analysis, and using the fourth sensor surface as a reference. A multi-channel system was used to screen the larger library (FL1056) since it enabled screening with a high throughput (Fig. 2b, bottom). It allowed rapid addressing of 16 biosensor surfaces via parallel injection of analyte across 8 channels. It was also used for screening of FL90 against the FPPS variants, where each variant was immobilized in a separate channel and referenced by a blank surface for each surface. The high-throughput assay format was successful in identification of hits for all targets (Fig. 1).

### Library pre-screening

Pre-screening was performed when using FL1056 but not FL90 since suitable screening conditions had already been identified for FPPS and FL90.^18^ The advantage of pre-screening is illustrated with a representative dataset for FL1056 and AChBP (Fig. 4a). Troublesome fragments were identified by distorted signals with a high response and trailing signals in subsequent injections (Fig. 4a, left). By removing the fragments, a clean data set was obtained for the screen (Fig. 4a, right). The pre-screening routine resulted in omitting approximately 1% of the compounds, depending on the experiment/target, *i.e*. on the experimental conditions used in each case (Fig. 1).

**Fig. 4.**
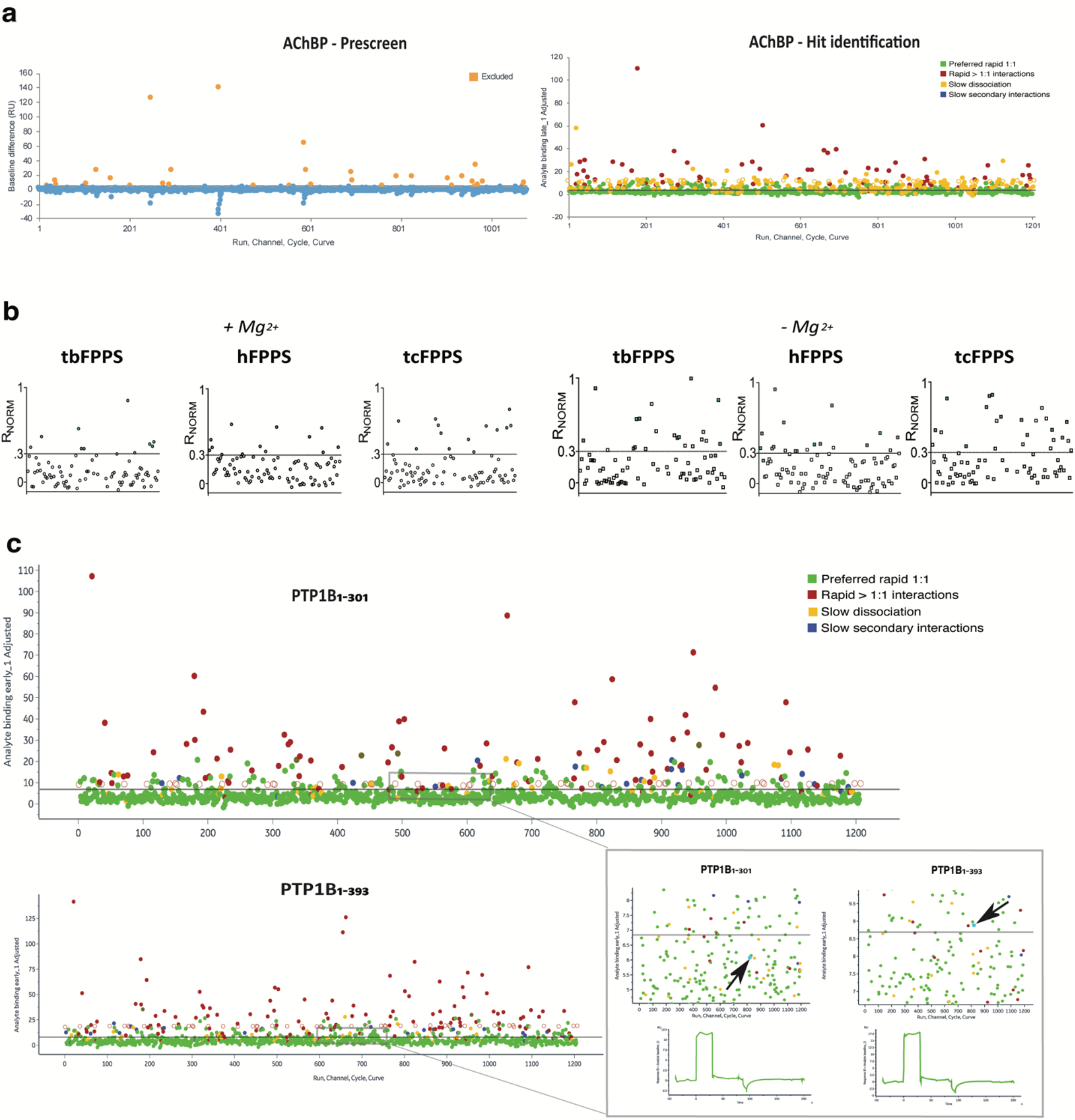
Fragment library pre-screening, screening and hit identification. Fragments were injected over sensor surfaces at concentrations and in running buffers specified in Fig. 1a. Screening with the single channel system used a 60 s contact time, 60 s dissociation time and a flow rate of 30 μL/min, while a 30 s contact time and 15 s of dissociation time was used with the multi-channel system. The flow system was washed with 50% DMSO after each cycle. **a,** Left, Data from pre-screening FL1056 against AChBP, identifying potentially troublesome compounds. Fragments were injected as in the actual screen (Fig. 1a, screening), but with a shorter cycle time (10 s contact time and 0 s dissociation). Fragments resulting in baseline changes between injections of at least 10 RU were subsequently excluded from screening experiments (see Fig. 1a). Right, Fragments interacting without discernible secondary binding complexities and response levels above a threshold set to result in the selection of 10% of the screened compounds are considered hits. The colour coding in the Hit identification plot is based on fragment kinetic profiles (Fig. 2 e): preferred rapid 1:1 (green), rapid interactions >1:1 (red), slow dissociation (yellow) and slow secondary interactions (blue). **b,** Data from screening FL90 against FPPS from *human, trypanosoma cruzi* and *brucei* in the presence (left) and absence (right) of the cofactor Mg^2+^. Fragments with normalised SPR signals (*R*_norm_) between 0.3 and 1 in the presence of Mg^2+^ were defined as hits (green). **c,** Example data sets from screening FL1056 against PTP1B_1-301_ (top) and PTP1B_1-393_ (bottom). A black arrow marks a fragment identified as a hit for one target but not the other, illustrating that automated data analysis can potentially mislead hit identification (inset).

### Example 1: Screening fragments against targets without tool compounds – FPPS

Farnesyl pyrophosphate synthase (FPPS) (See SI) was used to demonstrate that multiplexed approaches can be used to overcome the need for 1) tool compounds for checking target functionality during fragment screening, and 2) active site-binding ligands as competitors for identifying allosteric ligands. The first approach exploited the fact that FPPS is dependent on Mg^2+^ as a cofactor, structuring the active site in a catalytically competent state. By screening in the absence and presence of the cofactor Mg^2+^, and using the experiment without the cofactor as a negative reference, it was possible to identify fragments selective for the structurally intact protein (Fig. 4b).^18^ The reliability of this approach was supported by the observation that a larger number of fragments interacted with the surface in the absence of Mg^2+^, indicative of non-specific binding to unfolded regions.

The second approach involved screening towards FPPS from three different species. It does not rely on a reference surface with a blocked active site, but assumes that allosteric sites are not conserved between species and that potentially allosteric ligands can be identified among fragments that only interacted with one FPPS variant (*i.e*. not the conserved active site). This approach is suitable also for identifying species-selective ligands.

The data presented in this example is based on screening of a small library (FL90) in three independent experiments. Initially, a single channel system was used to address the three FPPS variants and a reference surface with the same injection of analyte (two independent screens). As an independent control, the library was also screened using a multi-channel system where each species variant of FPPS was immobilized in a separate channel also containing a blank reference surface.

Although the fragments showed suboptimal interaction kinetic profiles with secondary effects (*e.g*. Fig. 5d and e), the same hits were identified irrespective of the system, experimental design or immobilisation method, confirming the reliability of hit identification. The hit rate was ~15%, with at least 10 hits identified for each target (Fig. 1). Fragments previously identified to interact with tcFPPS were identified as hits also in these experiments.^18^ Due to very low affinities, saturation of the binding was not seen with the highest concentrations in hit validation step using concentration series (Fig. 5m-u). It was therefore not possible to estimate *K*_D_-values for the hits. Interactions were consequently quantified without a mechanistic/stoichiometric interpretation, via estimation of BE from the interactions at low concentrations (explained in Fig. 2g).

**Fig. 5.**
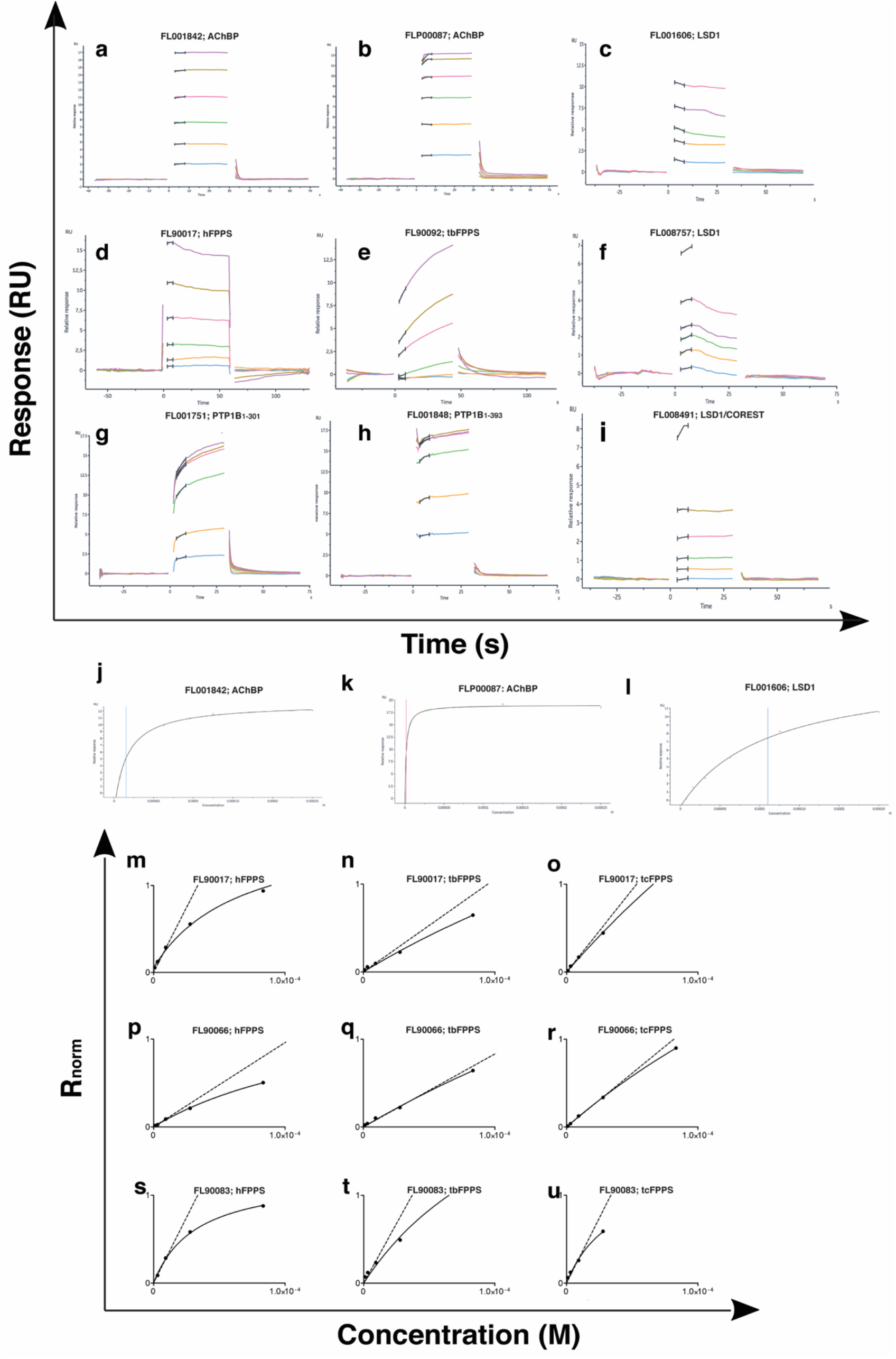
Secondary screening data and hit confirmation. FL90 hits were confirmed in a three-fold dilution series starting at 250 μM for 60 seconds at a flow rate of 50 μL/min, using a single channel system. FL1056 hits were confirmed in a two-fold dilution series starting at 250 or 500 μM, depending on the target (see Fig. 1) for 30 seconds at a flow rate of 30 μL/min., using a multi-channel system. The flow system was washed with 50% DMSO after each cycle. Zero concentration injections were used for blank subtraction. Non-specific signals were removed by subtraction of signal from reference channel. Solvent correction was performed with 8-point samples at appropriate DMSO concentrations. The dose response curves were fitted using a 1:1 binding model with free *R*_max_. **a-c,** Examples of sensorgrams with square pulse typical for fragments, *i.e*. fast association and dissociation kinetics (Fig. 2e, green). **d-i,** Examples of sensorgrams for fragments exhibiting non-ideal interactions (Fig. 2e, red, yellow, blue). **j**, signal vs. concentration curves for sensorgrams in a-c. **j-l**. Solid curves are based on *R*_norm_ data fitted by nonlinear regression analysis to a simple 1:1 interaction (insufficient for quantification due to *K*_D_≫ screening concentration). The dashed lines **m-u)** dose response plots for FPPS with the tangent represent the slopes of the graphs at low ligand concentrations, from which BE was estimated.

Three orthogonal methods were explored for confirmation of validated FFPS hits. Due to low affinities, none of these were ideal. 1) Microscale Thermophoresis (MST) showed weak concentration-dependent changes in thermophoresis (data for the three tcFPPS hits with the clearest results are shown in Fig. 6a). However, as for the SPR experiments, *K*_D_-values could not be quantified since saturation was not reached within the concentration range that could be used. 2) DSF confirmed the interactions, but resulted only in very small changes in T_m_ (Fig. 6d). 3) X-ray analysis of co-crystals of the hits with their respective targets could not confirm if fragments were present or not. However, the resolution was poor (the highest being only 2.6 Å) and, considering the weak affinities of the hits it is possible that the binding sites are only sparsely populated and potentially disordered.

**Fig. 6.**
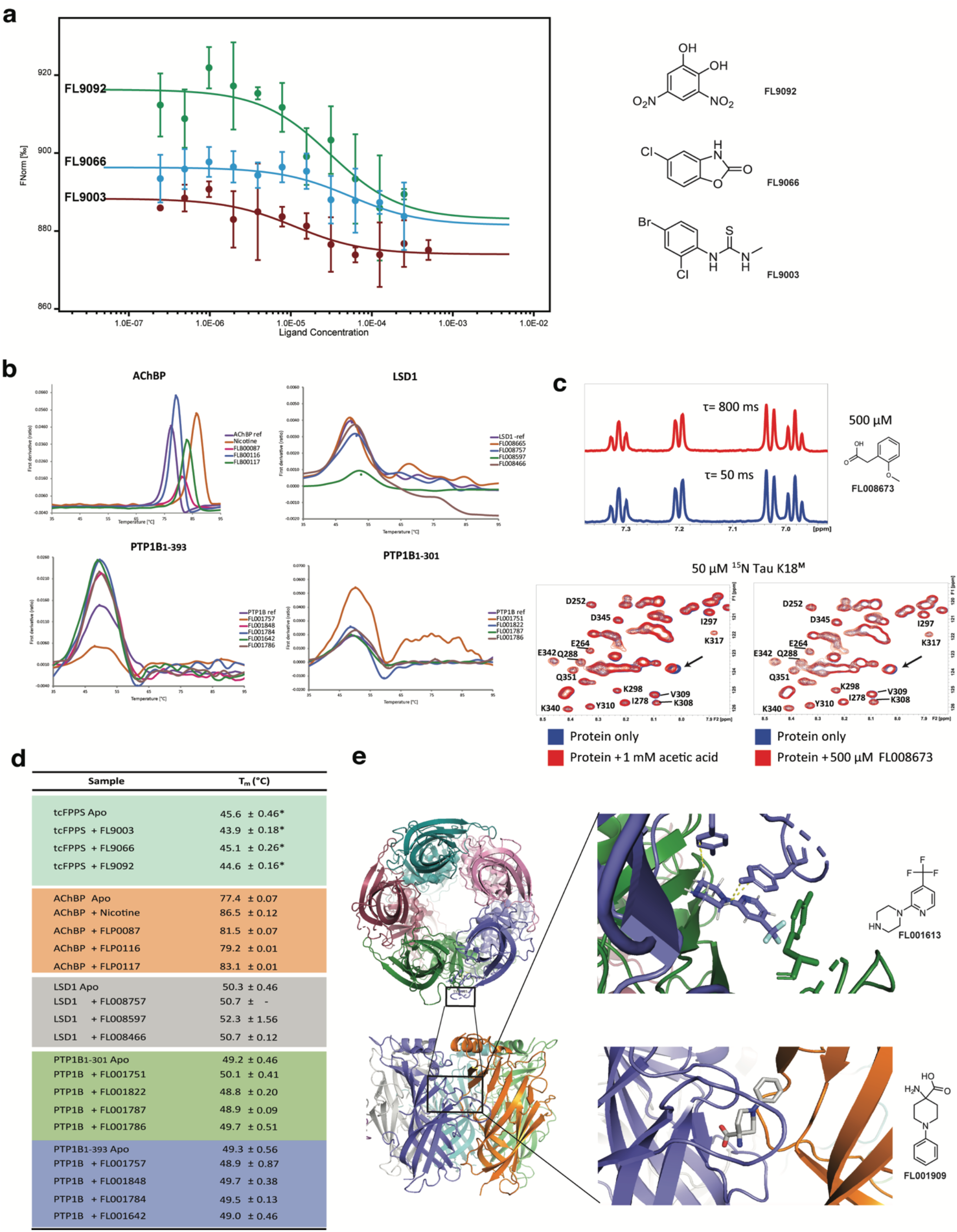
Validation of hits using orthogonal methods. **a,** MST traces for tcFPPS and identified hits. **b,** DSF traces for representative hits. **c,** 1D ^1^H CPMG-filtered NMR spectra at different delays for a selected fragment at 500 μM. Comparison of 2D ^1^H-^15^N SF-HMQC spectra of ^15^N tau K18^M^ in the presence of FL008673 (left) and acetic acid (right). **d**, T_m_ for hits determined by DSF, *is data from.^18^ **e,** Crystal structure of AChBP with fragment bound at Cys-loop interface between each monomer of pentameric structure.

The screening hits in this example thus appeared to have too low affinity for reliable confirmation by these orthogonal methods. Still, it demonstrates that SPR is the most sensitive of these methods. The next logical step in this project would be use the developed SPR biosensor assay to identify hits with higher affinities by screening a larger library or hit analogues.

### Example 2: Identification of fragments interacting with complex proteins – LSD1

Lysine demethylase 1 (LSD1) is a large and complex target with multiple functions (see SI). It has FAD as a catalytic cofactor and is regulated by interactions with COREST, a protein cofactor that stabilizes the active structure of the protein. Ligands targeting the protein substrate site, the FAD co-factor site, potential allosteric sites or the LSD1/COREST proteinprotein interaction surface, are all of interest since the ideal binding site or mode-of-action of ligands are not known. This project served as an example of how to use referencing that does not rely on reference compounds. It had access to a compound for monitoring immobilisation and functionality of LSD1 alone, but it does not interact with LSD1 in the presence of COREST. No tool compound was available for verifying the structural integrity of the LSD1/COREST complex.

By multiplexed screening of FL1056 against LSD1 alone and the LSD1/COREST complex, it was possible to discriminate hits interacting with different parts of the protein affected or not by the cofactor (Fig. 1a). Initial hit calling was established from assays using a high-density sensor surface while subsequent follow-up and verification used lower density surfaces. This permitted the detection of weakly binding fragments, whilst also controlling for limited mass transport and steric hindrance artifacts. In some cases, the data from higher concentrations of fragments had to be omitted due to strong non-specific binding events, as exemplified with LSD1 (Fig. 5f and i).

Hits were selected on the basis of sensorgram shape and affinity, estimated from dose dependencies representing micro molar *K*_D_-values. Interestingly, a larger number of hits were identified in the primary screen against LSD1 alone (#11), compared to that against LSD1/COREST (#5). This suggests that it could be a hotspot for fragment binding, and that it may be possible to discriminate hits specific for the planar surface of the tower domain, or that are not sensitive to the binding of the cofactor.

Orthogonal confirmation was limited to thermal shift analysis since crystallisation of fragment-LSD1 complexes in the absence of COREST has not been successful so far and NMR was not an option due the large size of the protein. Of the seven hits selected for thermal shift analysis, three showed a positive Tm shift between 0.7 - 2.9 °C, indicating a stabilising effect or compaction of protein, not seen in the DMSO control (Fig. 6b and d). One fragment showed a negative T_m_ shift and three were silent in the assay. Further confirmation of these hits and identification of their binding sites remains to be established. As for FPPS, the next logical step would be use the developed SPR biosensor assay to screen hit analogues for compounds with higher affinities. This would enable more elaborate competition experiments for identification of binding sites while interaction experiments using point mutations could confirm emerging binding hypotheses.

### Example 3: Identification of fragments interacting with structurally dynamic targets – AChBP

Acetylcholine binding protein (AChBP) was also used to illustrate the potential of SPR to identify fragments that interact with a large target, but for which the challenge is ligand induced conformational changes and multiple potential binding sites (See SI). AChBP is a proxy for the large class of therapeutically relevant human ligand gated ion channels (LGICs). It lacks the transmembrane domain, but provides a starting point for the identification of novel ligand scaffolds and binding sites. Still, its large size, pentameric structure with multiple known ligandable regulatory binding sites and a conformational flexibility remains a challenge.

To reduce these challenges, the target was immobilized using procedures optimised for orienting the pentamer on the surface. In addition, to avoid introducing methods for reliable selection of hits showing secondary effects, such fragments were not selected as hits, even when they gave high signals (Supplementary Fig. 5). The screening of FL1056 against AChBP showed that hits could readily be identified from single concentrations (Fig. 4a). Hits were validated by injecting the fragments in a concentration series. Since the interactions reached steady-state and the concentration range used was sufficient for saturation of the binding (Fig. 5j-l), *K*_D_ values were reliably estimated (Fig. 1c).

A set of ten AChBP hits were brought through to orthogonal confirmation using DSF and X-ray crystallography. All fragments resulted in a positive Tm shift between 2.5 - 9 °C in thermal shift studies (Fig. 6b and d), consistent with a stabilising effect or a structural compaction of the receptor, not seen by injecting DMSO alone. The binding sites and binding modes of the hits were established using X-ray crystallography. After numerous iterations of optimizing crystallography conditions, structures of 7 fragments in complex with AChBP were obtained with diffraction ~2.5 - 3.5Å. These structures readily identified fragment binding sites at the Cys-loop interface between monomers (Fig. 6f). The evolution of these fragments into leads for human LGICs can be guided via SPR-based interaction kinetic analysis using chimeric LGICs based on AChBP.^28^

### Example 4: Identification of fragments interacting with proteins containing intrinsically regions – PTP1B

Protein tyrosine phosphatase 1B (PTP1B) represents a target containing a C-terminal intrinsically disordered region (IDR). Here, multiplexed screening was performed against two truncated variants of PTP1B: PTP1B_1-301_ encompassing the folded catalytic phosphatase domain, and PTP1B_1-393_ also encompassing the intrinsically disordered region (Fig. 4c). It thus enabled the identification of two classes of hits, the first potentially specific for the IDR included in PTP1B_1-393_, and the second for the folded domain common to PTP1B_1-301_ and PTP1B_1-393_.

FL1056 Screening data for the two variants of PTP1B were normalized to account for differences in immobilization levels and functionality. Hits for the two surfaces were compared (Fig. 4c). To avoid potential minor mismatches between the surfaces, the standard automatic hit prioritization workflow was complemented by a manual assessment of the hits. The importance of a manual control of selected hits is illustrated for a hit automatically selected for PTP1B_1-393_, but not for PTP1B_1-301_ (Fig. 4c, bottom, left).

None of the fragments potentially specific for the IDR could be validated by the SPR assay when re-analysed and they were therefore not brought through to orthogonal confirmation. However, the fragments potentially interacting with the folded catalytic domain were confirmed using DSF. Of the six hits selected for PTP1B_1-301_, four exhibited T_m_ shifts ranging from 0.5 to 2.2 °C (Fig. 6b and d). Of those four fragments, two increased the T_m_ while the other two reduced the T_m_. For PTP1B_1-393_, four out of eight hits exhibited T_m_ shifts ranging from 0.5 to 1.9 °C. Three out of those four fragments increased the T_m_ while only one fragment caused decreased the T_m_ (Fig. 6d). Further confirmation of the binding sites and binding modes remains to be done. The developed SPR assay for these target variants is expected to be useful also for the initial stage of lead discovery since X-ray crystallography may be futile for these IDR containing proteins.^29^

### Example 5: Identification of fragments interacting with aggregation-prone proteins – K18^M^

An engineered monomeric form of tau K18 (tau K18^M^) was exploited as a model system for an aggregation-prone target (See SI).^30^ Two surfaces were used to explore the importance of orienting the monomeric form of the target on the surface. Screening of FL1056 was performed against tau K18^M^ immobilized using amine coupling, a method resulting in a heterogenous surface with random orientation of the target (SI figure 4). The identified hits were then re-analysed against biotinylated Avi-tagged tau K18^M^ immobilized via streptavidin capture, thus fixing the orientation of the protein on the surface. The outcome was different, indicating that the mode of immobilisation of the protein was important when selecting hits.

Orthogonal conformation of selected hits was done using NMR and ^15^N-labelled tau K18^M^ (Fig. 6c) since the simpler methods (DSF and MST) used for the other targets were not practicable for this target. However, 2D ^1^H-^15^N SF-HMQC NMR did not provide evidence of target-specific interactions with any of the selected fragments. A first control experiment, using 1D ligand-observed ^1^H CPMG-filtered NMR^31^ of the fragments, showed that this was not a consequence of aggregation at concentrations used during the analysis. A second control experiment revealed that the observed CSPs in the 2D experiments were similar to the ones produced in the presence of acetic acid, indicating that the fragments interacted non-specifically. Since the majority of hits (>90%) contained negatively charged groups, *e.g*. carboxylates, it can be speculated these interact with the highly positively charged tau K18^M^ surface at pH 7.4. The next step in this project would be to modify the developed SPR assay for screening under conditions where non-specific electrostatic interactions are minimised (*e.g*. optimising running buffers with respect to pH and ionic strength), and to avoid fragments with negative charges.

### Overall outcome of screening experiments

In total, approximately 107-110 fragments were identified as hits (Fig. 1). It follows from carrying through of an average of 10% from the single concentration screen to hit a concentration series validation in each screening campaign. Suboptimal interaction profiles were common in the FL90 screen against FPPS, most likely due to the features of the targets rather than the library. Simple interaction profiles, *i.e*. rapid association and dissociation kinetics and expected binding stoichiometry, were typically observed in the FL1056 screens, and most commonly for AChBP. In some cases, the data from higher concentrations of fragments had to be omitted due to strong non-specific binding, as exemplified with LSD1. Steady state was not achieved within the injection time for several hits, particularly for the IDPs. A pseudo steady-state analysis was useful as a means of establishing a concentration dependency, but not for quantification of affinities.

Between 4–19 fragment hits (*ca*. 1 to 10% of the screened library) were selected for orthogonal hit confirmation, depending on the target (Fig. 1). Methods with different experimental and detection principles were chosen specifically for each target. They were applied in order of how easily and fast results could be generated, *i.e*. typically first DSF or MST, then followed by X-ray crystallography or NMR. The success in this step varied, but was not pursued to the stage of giving conclusive results since it was not the focus of this study.

## Discussion

SPR biosensors and methods for identifying fragment hits have developed significantly since our first report of using this technology for FBDD.^20^ We have here provided new multiplexed methods for screening of a fragment library and identifying hits using contemporary SPR biosensors, thereby expanding the range of targets and libraries that can be used. The focus has been to illustrate experimental designs suitable with respect to the features of the target, desired hits, availability of target variants and tool compounds. The panel of targets was selected to illustrate different experimental challenges and for which the orthogonal conformation of hits therefore often was elusive, but outside the scope of the project. Importantly, alternative opportunities of pursuing lead identification using an SPR-based approach are highlighted.

A multiplexed strategy was developed for identifying potentially allosteric fragments, using different structural states of three species variants of the target (FPPS). It overcomes the requirement of active site binding tool compounds, otherwise enabling the screening against the target with a blocked active site which can directly identify allosteric ligands (Ref. Talibov et al, in revision).

Another challenge that can be overcome using a similar strategy involves screening against a target (LSD1) in the presence and absence of a protein binding partner (COREST), with the aim of identifying fragments with a potential to be evolved into leads perturbing protein-protein interactions. Since fragments are unlikely to have high enough affinities to interfere with a PPI, hits need to be evolved into more potent competitors before functional effects can be detected. By using an SPR-based, multiplexed approach, it is possible to guide optimisation without relying on functional or structure-based studies. Instead, experiments with either truncated or mutant versions of the protein binding partner can be useful for identifying the binding site.

For structurally dynamic targets, it is not only the location and structural features of the binding site that is of an aspect to consider, but also inherent challenges relating to the dynamics as such since energy losses arising from conformational transitions in the binding site affect the possibility to identify very weak interactions.^32^ Moreover, secondary effects resulting in distorted sensorgram can be confounding when selecting hits. In the example with AChBP, such fragments were not selected for validation and orthogonal confirmation, for simplicity. But they have been selected and analyzed in a separate study, and confirmed to induce conformational changes in the target (FitzGerald and Danielson, submitted manuscript). The possibility of simultaneously detecting binding and ligand-induced conformational changes using SPR biosensor assays^27^ enables the discrimination of fragments that simply bind to the target from those that also induce a conformational change.

The challenges of identifying ligands for dynamic target proteins is particularly difficult for intrinsically disordered proteins (IDPs), a class of drug targets that is not generally amenable to rational drug discovery methods. However, screening against an IDP is achievable if it has partially folded regions or if target variants can be engineered and tested in parallel. Several fragment hits were thus identified for the folded domain of PTP1B, but none of the hits potentially interacting specifically with the IDR of PTP1B could be confirmed, consistent with the elusive nature of unstructured regions.

The screening against tau K18^M^ resulted in a similar outcome, but also showed that there is a higher risk of detecting fragments interacting non-specifically with IDRs than with fully folded proteins (incidentally also seen with FPPS). Such effects can potentially be counteracted by optimising the conditions, *e.g*. using higher salt concentrations. However, it is difficult to optimise the experimental conditions for an IDP as it requires a good understanding of the structural and physico-chemical characteristics of the protein. It was shown also for tau K18^M^ that the immobilisation strategy can be critical. Procedures immobilising the target to the surface via a single attachment point, *e.g*. using biotin-streptavidin or antibody capture, can be used to avoid immobilising the target in a nonfunctional or non-native conformation, as might be the case when using a multipoint attachment, such as amine coupling. Engineering of the target for optimal immobilisation may therefore be beneficial.

## Conclusions

A broad range of targets can be used for SPR-driven FBDD, but their characteristics, availability of tool compounds and orthogonal assays for hit confirmation influence the chance for success. Practical solutions to challenging targets are emerging and they do therefore not have to be seen as inherently problematic but simply require additional assay development. The identification of fragment hits needs to consider the weak signals, rapid kinetics and low affinities expected from fragments. For success, it is critical to use high quality protein that can be immobilised to sensor surfaces in a fully functional form and at a high density. Although target variants or tool compounds are not essential, they increase the experimental repertoire and reliability of screening considerably.

The commercial availability of small fragment libraries, for example designed for crystallography studies, provide a useful option for academic settings. They allow SPR biosensors not designed specifically for fragments ssstudies to be used, and reduce solubility problems, handling issues and experimental time, while allowing chemical space to be adequately explored. However, recent technology developments enable higher throughput and faster screening campaigns, which benefits the screening of larger libraries. The highest throughput can be achieved using systems with single sensor surfaces in parallel flow channels, but limits the criteria for selection and prioritization of hits and results in a higher consumption of proteins and fragments. A more elaborate experimental design can be achieved using flow cells with several sensor surfaces, allowing the identification of unique hits based on multiple selection criteria and lower material consumption, but lower throughput. There are numerous ways to pick hits, and the method should be selected with respect to several criteria, including the experimental repertoire for orthogonal confirmation available in the lab. By multiplexing assays, some challenges of targets considered challenging for an SPR-based strategy can be overcome.

## Acknowledgments

The authors wish to acknowledge support from Eldar Abdurakhmanov and Annette Roos, SciLifeLab Drug Discovery and Development Platform, and library access from the Chemical Biology Consortium Sweden (CBCS). To members of the Danielson Lab for helpful discussions, to Olof Karlsson and the entire team at Cytiva for their continued support with this project. Furthermore, we wish to acknowledge colleagues from Beactica, Matthis Geitmann and Johan Winqvist for insightful discussions on fragment screening. The authors also wish to acknowledge Prof. Chris Ulens, Laboratory of Structural Neurobiology, KU Leuven for AChBP expression plasmids. Prof. Yang Shi and Benoit Laurent, Harvard Medical School. for LSD1/COREST plasmids. This project has received funding from the European Union’s Framework Programme for Research and Innovation Horizon 2020 (2014-2020) under the Marie-Skoldowska-Curie grant agreement numbers ID 675899 Fragment based drug discovery Network (FRAGNET) and Accelerated Early stage drug discovery (AEGIS) grant agreement ID 675555.

## Author contributions

U.H.D. conceptualized and supervised the overall project. I.J.P.E., M.W., P.O.B., H.F.K., J.E.M.K. and D.J.H. provided synthetic compounds. E.A.F. collated and curated fragment library FL1056. H.F.K. performed library analysis. E.A.F., V.O.T. and M.A. produced AChBP, LSD1 and LSD1/COREST. E.A.F. designed and performed SPR experiments with the proteins. E.A.F., P.B., D.E. and D.D. did crystallographic studies with AChBP. D.V. produced PTP1B and tau K18^M^ variants, and designed and performed SPR and DSF experiments with the proteins. D.V. performed NMR experiments with tau K18^M^. G.O. produced FPPS variants, and designed and performed SPR, DSF and MST experiments with the proteins. E.A.F., D.V., and G.O. drafted the manuscript. E.A.F. and D.V. prepared figures. E.A.F. and U.H.D. finalized the manuscript.

## Methods

### Protein production

#### Farnesyl pyrophosphate synthase (FPPS)

Production of N-terminally His-tagged farnesyl pyrophosphate synthase from *Trypanosoma cruzi* (tcFPPS), *Trypanosoma brucei* (tbFPPS) and *human* (hFPPS) were carried out as previously described for tcFPPS.^18^ Proteins were concentrated via centrifugation using a filter with a 30 kDa cut-off (Amicon Ultra-15), and the buffer was exchanged to storage buffer using PD10 columns (Cytiva). The storage buffer was 25 mM HEPES, 5 mM MgCl2 and 1 mM TCEP pH 6.5, supplemented with 100 mM NaCl for hFPPS, 25 mM NaCl for tbFPPS and 200 mM NaCl for tcFPPS. Aliquots of the enzymes were flash-frozen in liquid nitrogen and stored at −80 °C. Protein purity was estimated by SDS-PAGE and the concentration by NanoDrop ND-1000 Spectrophotometer (Marshall Scientific).

#### LSD1 and LSD1/COREST

LSD1 and LSD1/COREST were expressed and purified as previously described.^26^

#### Acetyl choline binding protein (AChBP)

The *Spodoptera frugiperda* insect cell line (Sf9) was utilized for expression of His-tagged *Lymnaea stagnalis* acetyl choline binding protein (Ls-AChBP) by infection with pre-isolated baculoviral stock (passage five, P5) with pFastBac1 and Ls-AChBP gene fused in the viral genome, as described previously.^33^ The cells were grown in supplemented Insect-XPRESS (Lonza) (penicillin and streptomycin; 100 u/mL) at a cell density of 2 x106 cells/mL. 1 mL per 100 mL cell culture of P5 viral stock was added to initiate protein expression. The cells were left to incubate for 48 hours at 27 °C with 90 revolutions per minutes (rpm) in a Minitron incubator Shaker (Infors HT).

Infected cells were centrifuged for 20 minutes at 4000 rpm in an Avanti J-26S XP (Beckman Coulter) and the supernatant was then decanted into a separate flask. Ni Sepharose™ excel beads (Cytiva) were prepared by rinsing the beads in a wash buffer (20 mM Tris-HCl pH 8.0, 300 mM NaCl). 1 mL of pre-rinsed beads were added to 1 L of supernatant and left for stirring at a minimal speed for one hour at 4 °C. Next the beads were collected by filtrating the media with a filter funnel, the beads were then transferred to a PD-10column. The column was rinsed with an imidazole containing washing buffer (20 mM Tris-HCl, 40 mM imidazole pH 8.0, 300 mM NaCl) for three column volumes. The protein was then eluted with an elution buffer (20 mM Tris-HCl, 300 mM imidazole pH 8.0, 300 mM NaCl) and fractions were collected, and the protein concentration was measured with absorbance on ND-1000 spectrophotometer (NanoDrop^®^). The fractions containing protein were combined and a buffer exchange and the protein were concentrated with a 30 K Amicon^®^ Ultra Centrifugal Filter spin column (Merck KGaA) to the storage buffer (20 mM HEPES pH 7.4, 137 mM NaCl and 2.7 mM KCl). The protein concentration was additionally measured on ND-1000 spectrophotometer (NanoDrop^®^) and the protein quality was evaluated by nanoDSF using a Tycho (Nanotemper).

#### Protein tyrosine phosphatase 1B (PTPIB_1-301_)

The gene for the catalytic domain (residues 1-ca. 301) of human protein tyrosine phosphatase 1B (PTP1B) with an N-terminal 6xHis tag (PTP1B1-301) was kindly provided by N. Tonks (Cold Spring Harbour, USA) in a pRP1B expression vector. Expression was performed using a published protocol.^34^ In short, the plasmid was transformed into *E. coli* BL21 (DE3) RIL cells (Agilent) and expression was induced with 1 mM IPTG for 20 h at 18 °C with continuous agitation at 200 RPM in LB media. After expression, cell pellets were spun down and stored at −80 °C until further use.

For purification, cell paste was thawed on ice and resuspended in ice-cold lysis buffer (1:3 ratio w/v of cell paste to lysis buffer) (50 mM Tris-HCl, pH 8.0, 500 mM NaCl, 5 mM imidazole, 0.5 mM MTG). One tablet of Roche cOmplete protease inhibitor and 500 μL of 2 mg/mL DNAseI (Sigma) was added per 50 mL of lysate. The mixture was stirred at 4 °C for 30 min. The cells were then homogenized using a Stansted French press system and the cell lysate was clarified by centrifugation at 40 000 g with a JA 25.50 rotor for 40 min at 4 °C. The clarified supernatant was then loaded onto a 5 mL HisTrap FF column (Cytiva), equilibrated with Buffer A (50 mM Tris-HCl pH 7.5, 500 mM NaCl, 0.5 mM MTG). PTP1B1-301 was eluted using three-step gradient to Buffer B (5, 20, 100%) (50 mM Tris-HCl pH 7.5, 500 mM NaCl, 500 mM imidazole, 0.5 mM MTG). Fractions with acceptable purity were desalted to Buffer C (50 mM Tris-HCl pH 7.5, 25 mM NaCl, 0.5 mM MTG) using HiPrep Desalt 26/10 column (Cytiva) at 6 mL/min and loaded on pre-equilibrated with Buffer C anion exchange column Mono Q 10/100 GL (Cytiva) at 4 mL/min. PTP1B1-301 was eluted at 2 mL/min using gradient over 20 CV from 0 to 50% of Buffer D (50 mM Tris-HCl pH 7.5, 1 M NaCl, 0.5 mM MTG). Fractions containing the protein of interest were pooled, concentrated with Amicon 10K MWCO concentrator to 15 mL and loaded 3 times in total as 5 mL loads on HiLoad 16/60 Superdex 75 pg column (Cytiva) at 1 mL/min, pre-equilibrated with Buffer E (50 mM HEPES pH 6.8, 150 mM NaCl, 0.5 mM TCEP). Pure fractions were pooled, concentrated with Amicon 10K MWCO concentrator before assessing protein concentration using A280 with ε = 46940 1/M*cm. Protein samples were aliquoted, snap frozen in liquid N2 and stored at −80 °C. The intact mass was checked using HPLC-MS assay.

#### Protein tyrosine phosphatase 1B (PTPIB_1-393_)

A human PTP1B construct encompassing the N-terminal catalytic domain and the disordered region, with an N-terminal GST tag and a C-terminal 6xHis tag (PTP1B1-393), in a pET-24a expression vector was ordered from DNA 2.0 (USA). As for PTP1B1-301, the plasmid was transformed into *E. coli* BL21 (DE3) cells (Agilent) and the expression of PTP1B1-393 was induced with 1 mM IPTG for 18 h at 18 °C with continuous agitation at 200 RPM in TB media. After expression, cell pellets were spun down and stored at −80 °C until further use.

For purification, cell paste was thawed on ice and resuspended to a 1:3 ratio in ice cold Lysis Buffer (50 mM Tris-HCl pH 8.0, 500 mM NaCl, 10 mM Imidazole, 0.1% Triton X-100, 0.5 mM DTT). One tablet of Roche cOmplete protease inhibitor and 500 μL of 2 mg/mL DNAseI (Sigma) was added per 50 mL of lysate. The mixture was left to stir at 4 °C for 30 min. The cells were then homogenized using a Stansted French press system. Following this, the cell lysate was clarified by centrifugation at 40 000 g with JA 25.50 rotor for 40 min at 4 °C. The clarified supernatant was loaded onto a 5 mL HisTrap FF column (Cytiva) twice, equilibrated with Buffer A (25 mM Tris-HCl pH 8.0, 500 mM NaCl, 0.5 mM DTT). PTP1B_1-393_ was eluted using a three-step gradient (5, 20, 100%) to Buffer B (25 mM Tris-HCl pH 8.0, 500 mM NaCl, 500 mM imidazole, 0.5 mM DTT). Fractions with acceptable purity were desalted to Buffer C (25 mM Tris-HCl pH 7.4, 150 mM NaCl, 1 mM EDTA, 1 mM DTT) using HiPrep Desalt 26/10 column (Cytiva) at 6 mL/min, whereafter 150 μL of PreScission protease (Cytiva) was added to the mixture for overnight incubation at 4 °C. After QC with SDS-PAGE to see if the cleavage reaction is complete, the sample was loaded onto preequilibrated with Buffer C 5 mL GSTrap HP column (Cytiva) at 1 mL/min. Cleaved PTP1B_1-393_ was found in flow-through fractions whereas contaminants were found in the 100% elution fractions with Buffer D (25 mM Tris HCl pH 8.0, 150 mM NaCl, 10 mM reduced glutathione, 1 mM DTT). Following this, PTP1B_1-393_ was desalted to Buffer E (25 mM Tris-HCl pH 7.5, 25 mM NaCl, 1 mM DTT) using HiPrep Desalt 26/10 (Cytiva) at 6 mL/min and loaded on a Mono Q 10/100 GL column (Cytiva) at 4 mL/min. The protein was eluted using gradient from 0 to 30% over 12 column volumes with Buffer F (25 mM Tris-HCl pH 7.5, 1 M NaCl, 1 mM DTT). Pure fractions were concentrated using Amicon 10K MWCO concentrator to 10 mL and further purified with pre-equilibrated with Buffer F (25 mM HEPES pH 6.8, 150 mM NaCl, 2 mM DTT) HiLoad 26/60 75pg column (Cytiva) at 2 mL/min. Protein concentration was determined using A280 with ε = 53400 1/M*cm. Protein samples were aliquoted, snap frozen in liquid N2 and stored at −80 °C. The intact mass was checked using HPLC-MS.

#### Tau K18^M^

An engineered human tau K18^M^ construct corresponding to the paired helical filaments binding domain (residues 244-372), with C291S and C322S substitutions keeping it as a stable monomer, was produced as previously described, including biotinylated and isotopically labelled constructs.^30^

### Surface Plasmon Resonance biosensor experiments

#### Biosensor surface preparations

##### FPPS

Three FPPS variants (hFPPS, tcFPPS and tbFPPS) were immobilized via amine coupling to CM5 sensor chips (Cytiva) at 25 °C and a flow rate of 10 μL/min, using standard procedures.^35^ For all three enzymes, the running buffer used for immobilization consisted of 10 mM HEPES, 150 mM NaCl, 3 mM MgCl2, 1 mM TCEP and 0.05% Tween-20. The surfaces were activated using a 1:1 mixture of 400 mM EDC+ 100 mM NHS for 210 s. The protein was injected at 50 μg/mL and 5 μL/min for a time resulting in an immobilization level of ~3000 to 5000 RU, generating a theoretical *R*_max_ of ~20 RU for a fragment molecule of ~150 Da. Unreacted carboxyl groups remaining on the surface were deactivated with 1 M ethanolamine chloride (pH 8.5) for 210 s.

##### AChBP

The surface of Sensor Chip NTA (Cytiva) was first conditioned with 1 min injection of regeneration solution 10 μL/min at 25 °C. This was followed by injection of running buffer to remove any excess regeneration solution. Next, the surface was prepared with 0.5 mM NiCl2 by injecting a 1 min pulse of nickel solution at 5 μL/min, with a subsequent wash step with running buffer supplemented with 3 mM EDTA. To enhance stability of the surface and to ensure correct orientation of ligand the sensor chip was activated using 1:1 mixture of 400 mM EDC+ 100 mM NHS for 420 s at 10 μL/min at 25 °C. This was followed by injection of 10 μg/mL target protein in its immobilization buffer for appropriate time at 10 μL/min until an immobilization level ~4000 RU corresponding to a theoretical *R*_max_ of 20-40 RU for a 150 Da fragment. Eight start-up cycles were used for stabilizing the surface after the immobilization.

##### LSD1 & LSD1/COREST

The immobilization methods were the same for Sensor Chip CM5 (Cytiva) and Sensor Chip CM7 (Cytiva). The surface of Sensor Chip CM7 (Cytiva) was activated using 1:1 mixture of 400 mM EDC/100 mM NHS for 420 s at 10 μL/min at 25 °C. This was followed by injection of 10 μg/mL target protein in its immobilization buffer for appropriate time at 5 μL/min. The surface was then deactivated with 1 M ethanolamine for 420 s at 10 μL/min. The reference flow cell was activated and deactivated using the same protocol but without protein.

##### PTP1B_1-301_/PTP1B_1-393_

The surface of a Sensor Chip CM5 (Cytiva) was activated using 1:1 mixture of 400 mM EDC/100 mM NHS for 420 s at 10 μL/min at 25 °C. This was followed by injection of 25 μg/mL target protein in 10 mM sodium acetate pH 5.5, 1 mM DTT at 10 μL/min to achieve *R*_max_ value of 20-40 RU for 150 Da fragment. The surface was then deactivated with 1 M ethanolamine for 420 s at 10 μL/min. The reference flow cell was activated and deactivated using the same protocol but without protein.

##### Tau K18^M^

The surface of a Sensor Chip CM5 (Cytiva) was activated using 1:1 mixture of 400 mM EDC/100 mM NHS for 420 s at 10 μL/min at 25 °C. This was followed by injection of 25 μg/mL target protein in 10 mM sodium borate pH 8.5 at 10 μL/min to achieve *R*_max_ value of 20-40 RU. The surface was then deactivated with 1 M ethanolamine for 420 s at 10 μL/min. The reference flow cell was activated and deactivated using the same protocol but without protein.

The surface of Sensor Chip SA (Cytiva) was first conditioned with three consecutive 1 min injections of 1 M NaCl + 50 mM NaOH at 10 μL/min at 25 °C. This was followed by injection of 100 nM of biotinylated protein in 50 mM HEPES pH 7.4, 150 mM NaCl, 0.05% Tween-20 at 5 μL/min to achieve theoretical *R*_max_ value of 20-40 RU for a 150 Da fragment. Eight start-up cycles were used for stabilizing the surface after the immobilization

#### Fragment library screening

##### FPPS

Screening of fragment library FL90 against three FPPS variants (hFPPS, tcFPPS and tbFPPS) was performed at 25 °C. For all three enzymes, the running buffer used for screening and interaction analysis consisted of 10 mM HEPES, 150 mM NaCl, 1 mM TCEP and 0.05% Tween20, supplemented with 1% DMSO. Screening was carried out in the presence and absence of 3 mM MgCl2 in the buffer to discriminate binding to functional and non-functional targets.

The complete library was screened using a Biacore 8K+ system (Cytiva). Fragments were injected for 60 seconds at a flow rate of 50 μL/min in a final concentration of 250 μM. Fragments with signals between 30% and 100% of a theoretical *R*_max_ were selected as hits. The analysis was based on the average report point signal 6 seconds after the beginning of the injection (binding early response) in order to compensate for secondary effects. Binding early values were normalized (*R*_norm_) with respect to the theoretical *R*_max_ of each fragment (as in Equation 4), in order to account for differences in molecular weight and protein immobilization levels.^21^

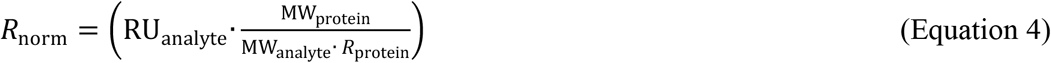

The selected hits were validated by analysis of the fragments in a 3-fold dilution series starting at 250 μM for 60 seconds at a flow rate of 50 μL/min on Biacore T200 (Cytiva).

##### PTP1B_1-301_ and PTPIB_1-393_

A comparative screening of FL1056 against PTP1B1-301 and PTP1B1-393 was performed using a Biacore 8K+ system (Cytiva). The proteins were immobilized on Sensor Chip CM5 (Cytiva) via amine coupling using 25 μg/mL protein in 10 mM sodium acetate pH 5.5, 1 mM DTT and aiming for 20-40 RU *R*_max_ for a 150 Da fragment. Suramin was injected as a positive control at 7.5 μM, the highest concentration that could be used without signals being affected by non-specific interactions (Supplementary Figure 3). Experiments were performed in 25 mM Tris-HCl pH 7.4, 150 mM NaCl, 2 mM EDTA, 1 mM DTT, 1% DMSO. Solvent correction was done with 0.5 to 1.8% DMSO. A hit threshold was set to 1 RU, based on blank controls, and 50% of normalized signals were used to distinguish IDR specific hits.

##### Tau K18^M^

6xHis-tau K18^M^ was immobilized on Sensor Chip CM5 (Cytiva) via amine coupling in 10 mM sodium borate pH 8.5 containing 25 μg/mL of protein to achieve *R*_max_ of 20-40 RU for a 150 Da fragment, followed by fragment screening campaign. Running buffer of 25 mM Tris-HCl pH 7.4, 150 mM NaCl, 2 mM EDTA, 1% DMSO was used for all fragment screening assays. Solvent correction was performed from 0.5 to 1.8%.

#### Interaction analysis

Fragments were injected over reference and immobilized protein surfaces at a single concentration (Fig. 1, Screening) in the appropriate running buffer for each target. Control compounds (Fig. 1, Assay design) were injected each 36^th^ cycle as a control for surface functionality over time (Supplementary Figure 1) DMSO solvent correction and reference surfaces were set-up in the same manner as for the single channel system. Experimental data was assessed using theoretically calculated *R*_max_ values instead of experimental *R*_max_ values since suitable tool compounds were lacking. In order to triage hits, a reductionist approach was taken and approximately the top 10% of hits, i.e. with the highest response and without undesired kinetics (slow association, slow dissociation or *R*_eq_≫*R*_max_, Fig. 2e) were prioritized.

Non-specific signals were removed by subtraction from reference channel, and solvent corrections were also performed to compensate for differences in DMSO concentrations. Apparent *K*_D_ values were estimated by steady state analysis using a 1:1 interaction model (Equation 2).

Hits were ranked on the basis of *K*_D_ or Binding Efficiency (BE) values, calculated as the initial slope of the linear relationship between complex concentration (in *R*_norm_) and ligand concentrations at very low ligand concentrations,^21^ The hit threshold was set to 30% of the theoretical *R*_max_ of each compound, with a limit at 100%. The data from all screens were evaluated with Biacore Insight Evaluation Software (Cytiva).

### NanoDSF analysis of target protein stability

The thermal stability of target proteins in different buffers and the potential stabilizing effects of ligands was evaluated at 1 μM using a direct thermal shift assay using a Tycho NT6 instrument (Nanotemper Technologies), monitoring intrinsic protein fluorescence at 330 and 350 nm during a thermal ramp from 35 °C to 95 °C. Data were plotted as a derivative to get the inflection point for the intrinsic fluorescence shift from which the inflection temperature (Ti) was determined.

The stabilizing effects of controls (where available) and hits from the FL1056 library screening against targets was analysed when possible. Samples were prepared by diluting protein and compound to a final concentration of 1 μM and 1 mM, respectively.

### Fragment library analysis

3-Dimensional structures were generated Pipeline Pilot 16.5.0.143, 2016, Accelrys Software Inc. Prior to conformer generation a wash step was performed, which involved stripping salts and ionising the molecule at pH 7.4. SMILES strings were converted to their canonical representation and the original stereochemistry at each chiral centre was recorded. Any stereo centre created during the ionisation would have undefined stereochemistry. A SMILES files was written that contained all possible stereoisomers of the molecule. Conformers were generated using Catalyst with the BEST conformational analysis method and relative stereochemistry. Catalyst was run directly on the server and not through the built-in Conformation Generator component. The maximum relative energy threshold was left at the default 20 kcal mol-1 and a maximum of 255 conformers were generated for each compound. The aim of this was to give the best possible coverage of conformational space. The resulting conformations from Catalyst were read and only those where the stereochemistry matched the original molecule or its enantiomer were kept. These were then all standardized to the original stereochemistry by mirroring the coordinates of the enantiomers. Duplicate conformations were filtered with a RMSD threshold of 0.1. Each conformation was minimized using 200 steps of Conjugate Gradient minimization with an RMS gradient tolerance of 0.1. This was performed using the CHARMm forcefield with Momany-Rone partial charge estimation and a Generalized Born implicit solvent model. After minimization, duplicates were filtered again with a RMSD threshold of 0.1.

Generated conformations were used to generate the three Principal Moments of Inertia (PMI) (I1, I2 and I3) which were then normalized by dividing the two lower values by the largest (I1/I3 and I2/I3) using Pipeline Pilot built-in components. PMI about the principal axes of a molecule were calculated according to the following rules: 1. The moments of inertia are computed for a series of straight lines through the centre of mass. 2. Distances are established along each line proportional to the reciprocal of the square root of I on either side of the centre of mass. The locus of these distances forms an ellipsoidal surface. The principal moments are associated with the principal axes of the ellipsoid.

Cumulative PMI analysis was performed in the following way. The ΣNPR (ΣNPR = NPR1 + NPR2) was calculated for each conformer and then the mean ΣNPR for each fragment was obtained. This value was used as a measure of the three-dimensionality of each fragment. The cumulative percentage of fragments within a defined distance from the rod-disc axis (ΣNPR) was calculated and plotted.

Molecular Weight (MW), heavy atom count (HAC), lipophilicity (SlogP), number of hydrogen bond donors (HBD), number of hydrogen bond acceptors (HBA), rotatable bond count (RBC), fraction of sp3 carbons (Fsp3) and topological polar surface area (TPSA) were calculated using RDKit v3.4 in KNIME v3.5.2. Prior to calculation, salts were stripped and canonical SMILES were generated. clogP values were calculated using Daylight/BioByte ClogP v4.3.

### Nuclear Magnetic Resonance

NMR data was collected using Bruker Avance III HD 600 MHz spectrometer with QCI cryoprobe at 298 K. The samples were prepared to contain 50 μM ^15^N-labelled tau K18^M^ in 25 mM Tris-HCl pH 7.4, 150 mM NaCl, 2 mM EDTA, 50 μM DSS, 1% d6-DMSO, 5% D2O.1D and 2D protein-observed NMR spectra were obtained using standard Bruker pulse sequences with the parameters specified in Supplementary Table 3. ^1^H-^15^N SF-HMQC was selected as it is more time-efficient with IDPs due to its lower spectral dispersion in 1H dimension in comparison to folded proteins.^36^

### Miscroscale Thermophoresis

Target proteins (hFPPS, tbFPPS, tcFPPS) were fluorescently labelled following the manufacturer protocol for free amine coupling of the dye (RED-NHS 2nd Generation (amine reactive)) (NanoTemper) to lysine residues. Stability of labelled protein was then verified using nanoDSF (see above “Target protein stability”), and their concentration by Nanodrop (DeNovix). MST analysis was carried out using premium-coated and standard treated capillaries (NanoTemper), and buffer matching the SPR conditions (HEPES 10 mM, NaCl 150 mM, MgCl2 3 mM, TCEP 1 mM, 0.05% Tween20) including 2% DMSO. Fragment dilutions were prepared by 1:1 serial dilution in 4% DMSO. Each dilution was then mix 1:1 with protein solution, to a final concentration of 500 - 0.244 μM fragment, and 562 nM target protein. Signals were recorded at MST 40%, Excitation 20% at 25 °C (Monolith NT.115)

### Co-crystallization of AChBP with compounds

AChBP at concentrations between 10 and 13 mg/mL in storage buffer (20 mM HEPES, 137 mM NaCl, 2.7 mM KCl, pH 7.4) was incubated with compound dissolved in DMSO, resulting in a final concentration of 2.5 mM compound and 5% DMSO. The drops of 2 μL contained a 1:1 ratio of protein-compound mix and reservoir solution (100 mM citric acid at pH 4.8-5.2 and 1.5-2 M ammonium sulphate).

The crystallization experiments, performed in a hanging drop vapour diffusion manner at RT, resulted in crystals of various morphologies forming after 1-2 weeks. The crystals were cryo-protected in a reservoir solution with 20% glycerol before snap-freezing.

Diffraction data was collected at Diamond Light Source (Oxford, UK) IO4 and MAXIV (Lund, Sweden) BioMAX beamline. Indexing, merging and scaling was doing using XDS, XSCALE, XDSCONVERT. Molecular replacement was done with PhaserMR^37^ with structure 1UW6^33^ as search model. The ligand dictionaries were created using AceDRG.^38^ Model building was done using Coot^39^ and structure refinement using REFMAC5.^40^ Figures were prepared with PyMol.

