## Supplementary information for "Multiplexed experimental strategies for fragment library screening using SPR biosensors"

This Supplementary Information includes Supplementary Figs. 1-5, Supplementary Notes 1 and 2, and Supplementary Tables 1-3.

### Supplementary Notes

#### Supplementary Note 1

##### *FPPS*

FPPS is a $Mg^{2+}$-dependent regulatory enzyme with a key role in the isoprenoid biosynthetic pathway and a target for osteoporosis and cancer therapy (hFPPS), as well as for new drugs against trypanosomiasis (tbFPPS, tcFPPS).[1] Although bisphosphonate substrate analogues are potential inhibitors for all FPPS isoforms, allosteric inhibitors have also been identified for the human isoform.[2,3] Allosteric ligands with novel scaffolds or binding sites are relevant for all three variants of FPPS since previously reported allosteric ligands have failed to exhibit anti-tumor activity in cell-based assays[3] and they have not yet been reported for tbFPPS or tcFPPS. The latter could result in antiparasitic drugs that do not target the human enzyme.

##### *LSD1*

LSD1 is an epigenetic target implicated in certain forms of cancer.[4] In many disease states including cancer and neurodegeneration, genes are over expressed, by targeting allosteric sites or indeed PPIs using novel biosensor-based methods may provide a unique opportunity to new therapeutic strategies. Histone methylation is an epigenetic modification that can repress or activate transcription of specific genes. Due to this activation or repression of genes studying these enzymes is of importance. In many diseases such as cancer, genes are over expressed. Targeting the cells expression machinery with activators or inhibitors may offer a unique therapeutic window and perspective to develop new drugs. Lysine specific demethylase 1 (LSD1) was first described in 2004 as the first histone tail demethylating enzyme.[5] This changed the previously thought permanent modification of methyl groups on lysine residues into a reversible process that regulates transcriptional levels.

.

##### *AChBP*

AChBP is a soluble homologue of human Cys-loop receptors, such as the nicotinic α7 receptor, which contains the extracellular ligand-binding domain but lacks the trans-membrane domain of this class of ligand-gated ion channels (LGICs). Notably, binding of ligands to AChBP induces the same conformational changes as in the human receptors, with a conserved activation mechanism. Although complete ion channels can be immobilised and used for screening by SPR, allowing the identification of ligands, the approach has so far only been demonstrated for recombinant homo pentamers.[6] AChBP is a simpler option for screening and orthogonal validation of hits, although for evolution of fragments into functional modulators, as agonists or antagonists, it is essential to use orthogonal assays that identify the binding site and effects of ligand binding on the structure for the actual target protein, or functional assays.[6–12]

##### *PTP1B*

Full-length PTP1B is a 434 aa protein with a folded N-terminal catalytic phosphatase domain (*ca.* 301 aa), a disordered proline-rich regulatory domain involved in protein-interactions (*ca.* 90 aa), and a hydrophobic C-terminal ER-targeting domain (*ca.* 40 aa). The protein is involved in diabetes, obesity development and mammary tumorigenesis.[13,14] The identification of IDPs has changed our understanding of the link between protein structure and function, and the importance of such interactions for regulation in mammalian biology.[15] There is consequently great interest exploring the possibilities of targeting and modulating IDPs with LMW ligands, with recent promising results.[16–19]

##### *Tau K18$^M$*

Human tau is an IDP involved in forming neurofibrillary tangles in Alzheimer´s disease. Tau K18 is a truncated form encompassing four paired helical filaments (PHFs) containing hexapeptide motifs at the core of formed fibrils. Fibril formation is enhanced by oxidation of Cys residues, why mutation of C291 and C322 to serine (C291S, C322S) can be used to generate a stable protein in the form of a monomer.[20]

#### Supplementary Note 2

##### *Resolution of complex interactions between suramin and PTP1B variants*

Suramin was confirmed to interact with $PTP1B_{1-301}$ and $PTP1B_{1-393}$ immobilised via amine coupling (Supplementary Figure 2).[21] The interaction was the same with both constructs, confirming that it interacts with the folded, catalytic domain of PTP1B and that it is not affected by the presence of the IDR region of $PTP1B_{1-393}$.[22] Nevertheless, PTP1B:suramin interactions are best described using a mechanistic model that accounts for binding to both a primary high affinity site and a secondary low affinity site (accounting also for additional lower affinity sites), as shown for $PTP1B_{1-321}$ (see Supplementary Figure 3). The $K_D$ values estimated for the PTP1B constructs (Supplementary Table 1) matched the reported values closely.

#### Supplementary Figures

**a**

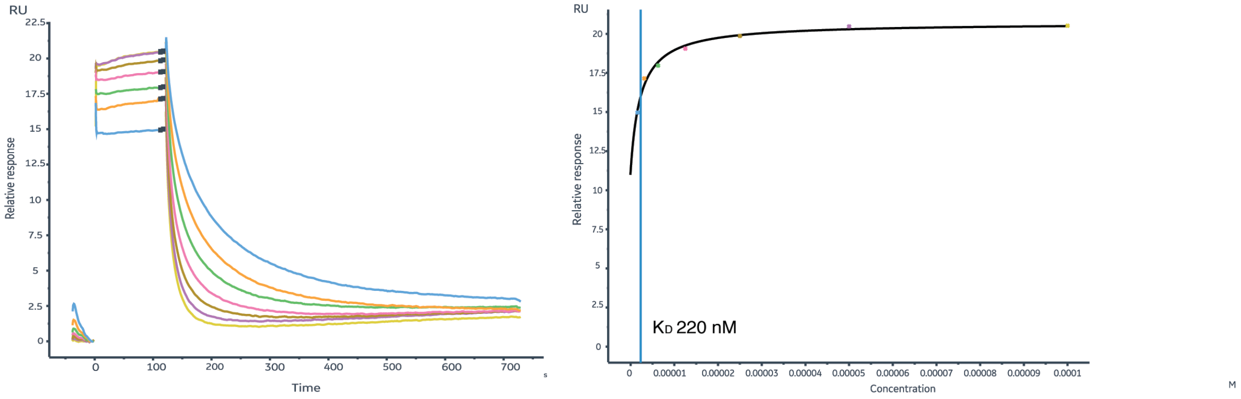


**b**

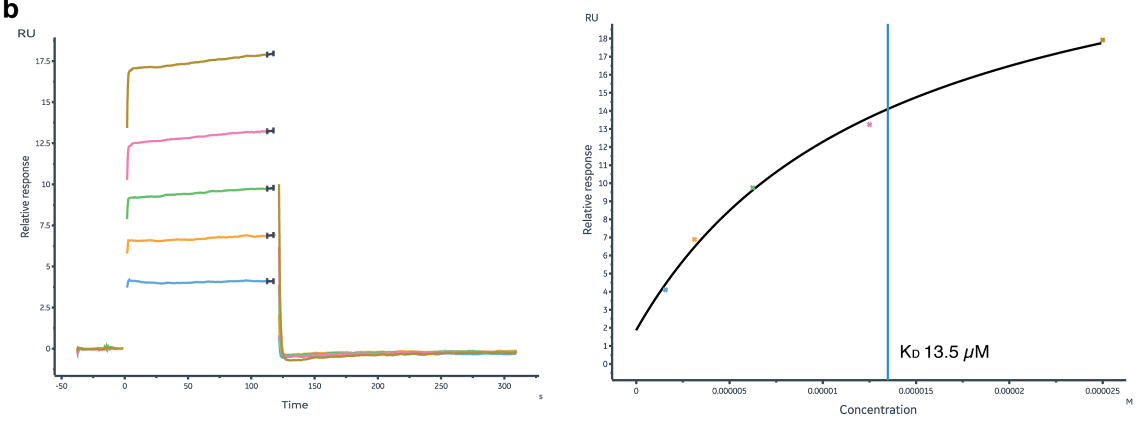


**Supplementary Figure 1. Sensor surface control experiments for AChBP and LSD1.** Sensorgrams and corresponding steady state plots for lobeline interactions with AChBP (a), and BEA1 interactions with LSD1 (b). The immobilization levels were ~3000 RU for AChBP and ~22000 RU for LSD1. The steady state plots are based on report points taken at the end of the injection (marked section). A 1:1 interaction mechanism was fitted to the steady state data by non-linear regression using the Biacore 8K software. The vertical line represents the ligand concentration equal to the estimated $K_D$-values. The same profiles were obtained when experiments were repeated after 48 hours *i.e.* the maximum time a surface was used in the screening campaign.

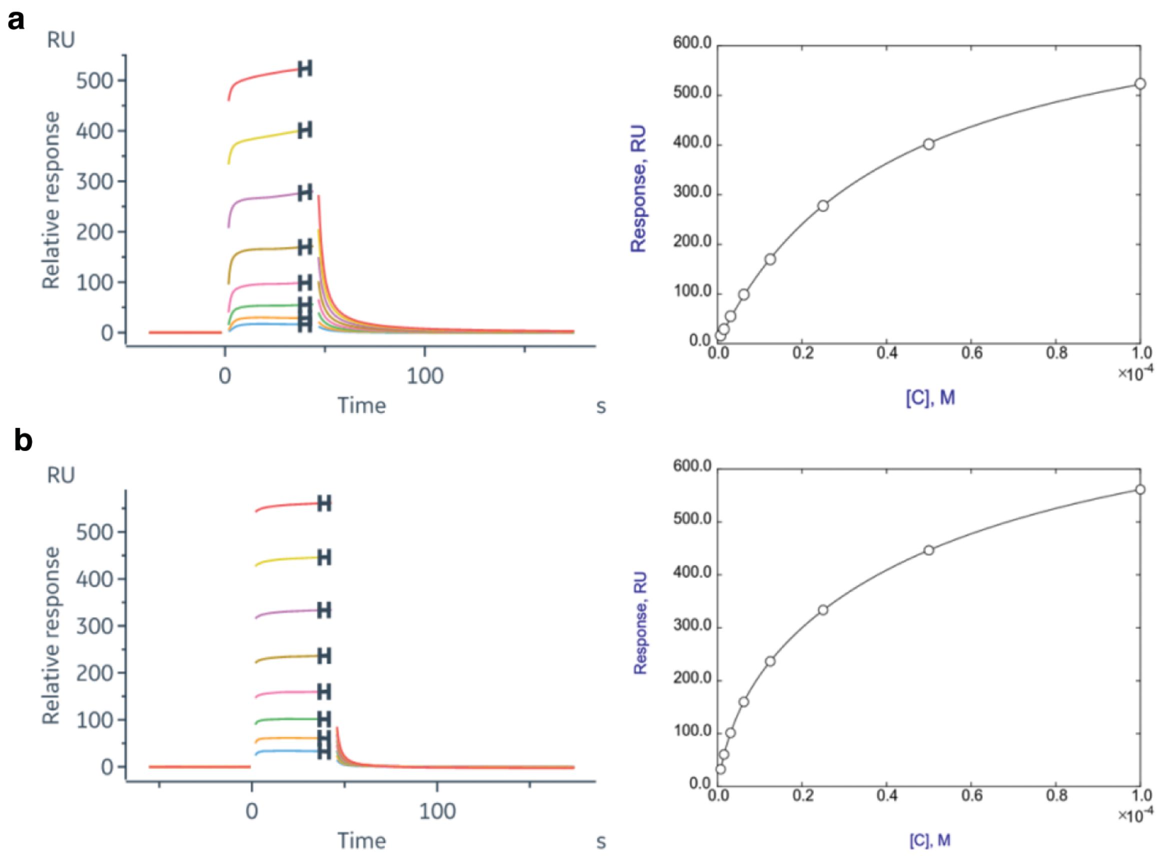


**Supplementary Figure 2**. **Sensor surface control experiments for PTP1B variants.** Sensorgrams and corresponding steady state plots for suramin interactions with PTP1B$_{1\text{-}301}$ (a), and PTP1B$_{1\text{-}393}$ (b). The immobilization levels were ~6400 RU for PTP1B$_{1\text{-}301}$, and ~9400 RU for PTP1B$_{1\text{-}393}$. The steady state plots are based on a two-fold dilution series ranging from 0.78 to 100 µM, and report points taken at the end of the injection (marked section). A 2:1 stoichiometry with a high and a low affinity site (See Supplementary Note 2) was fitted to the steady state data by non-linear regression, using SimFit (https://simfit.org.uk). The $K_D$ values are presented in Supplementary Table 1.

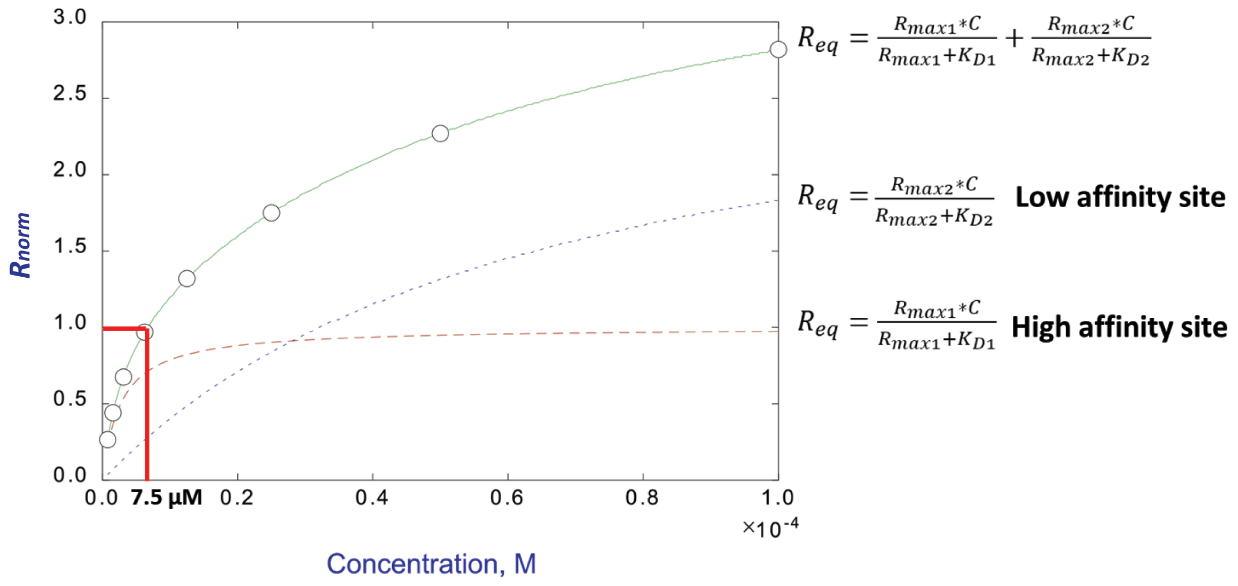


**Supplementary Figure 3**. **Differentiating high and low affinity interactions for suramin and PTP1B.** SPR signals for the interaction between immobilized PTP1B$_{1\text{-}193}$ and a concentration series of suramin at steady state (from Supplementary Figure 2b) was normalized ($R_{norm}$=Relative response·$MW_{protein}/MW_{analyte}\cdot R_{protein}$) with respect to the immobilization level of the protein ($R_{protein}$) and the molecular weights of the analyte and protein. The fitted curve represents a mechanistic model with two binding sites (solid line). The theoretical curve for the high affinity site (dashed) has $R_{max}$ = 1 at saturation, while the curve for the low affinity site (dotted) does not saturate at 2 or 3, indicating that it is composed of multiple lower affinity sites, typical of non-specific interactions. The non-linear regression analysis was carried out using SimFit (https://simfit.org.uk). The analysis was used to identify a suitable suramin concentration for control injections monitoring PTP1B sensor surface quality. A concentration of 7.5 µM (red line) was selected since it saturates the high affinity site ($R_{norm}$=1), giving a relatively high signal, without significant signals from low affinity sites.

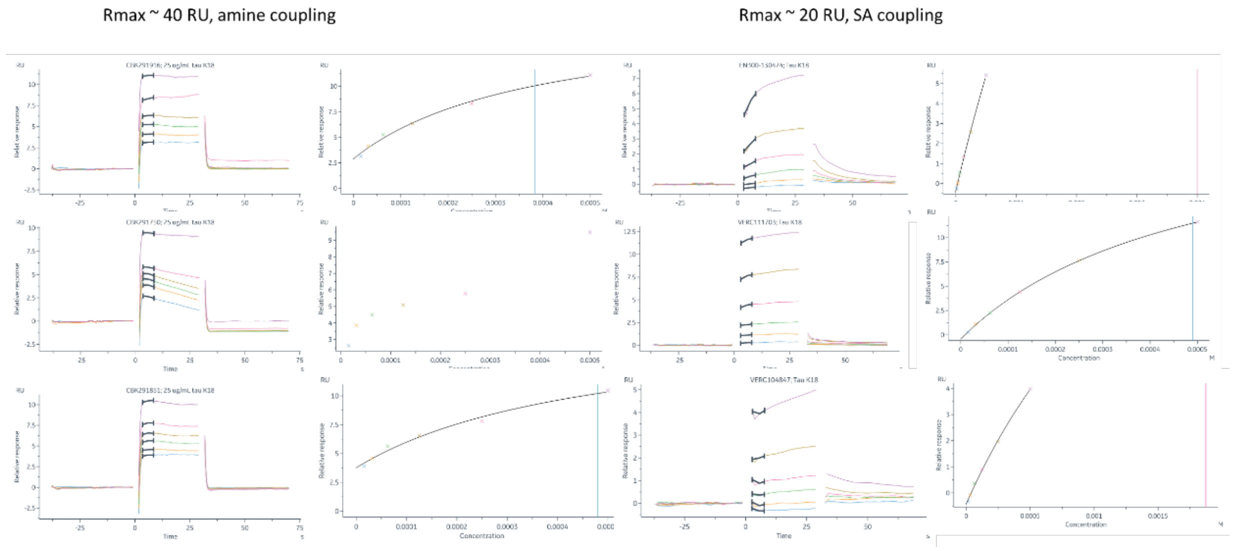


**Supplementary Figure 4.** Illustration of importance of using a suitable immobilization method. Sensorgrams and dose-response graphs for the same three fragments interacting with tau K18$^M$ immobilized randomly via amine coupling (**Left**) and oriented using biotinylated avi-tagged tau K18$^M$ captured via streptavidin (**Right**).

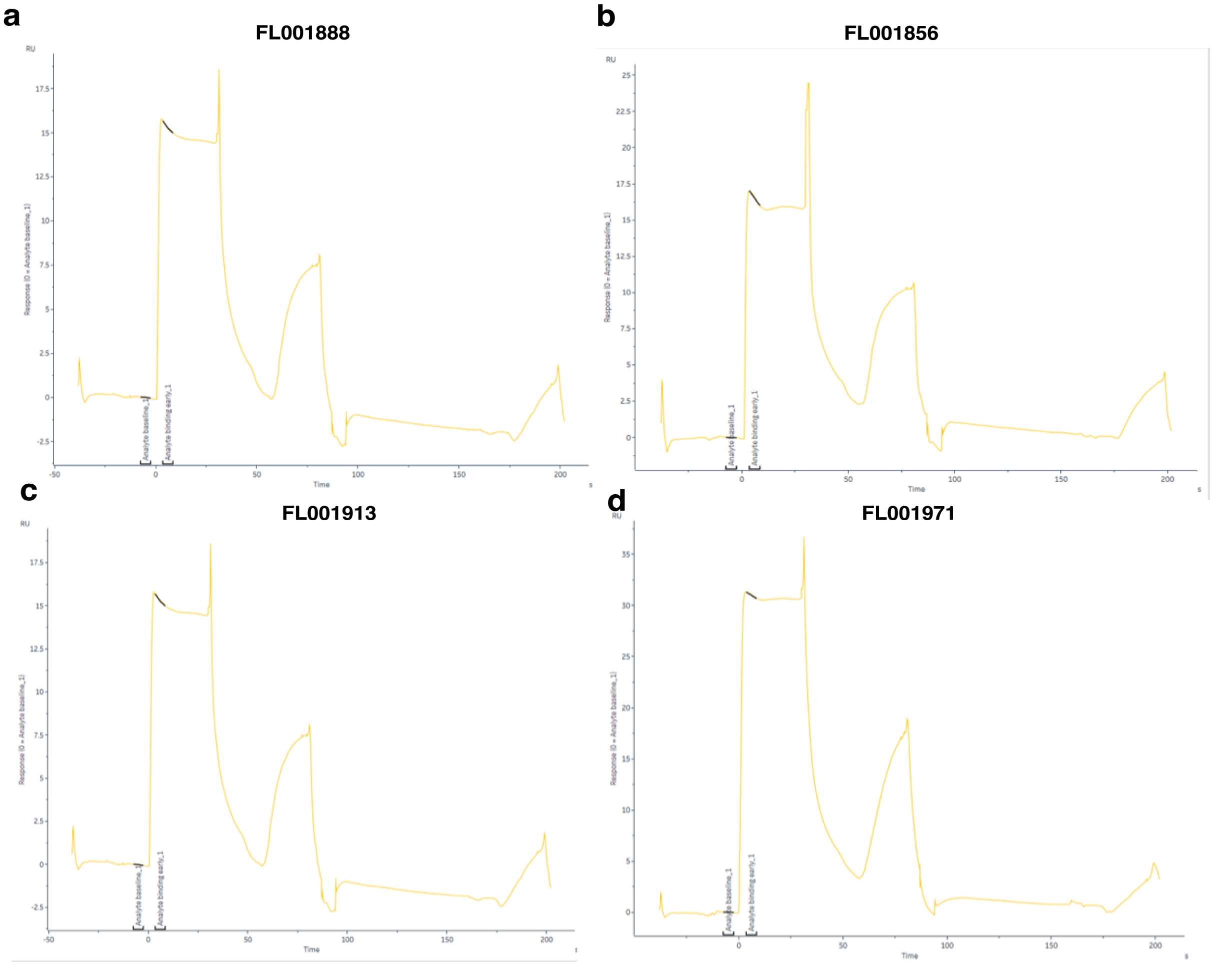


**Supplementary Figure 5**. **Risk of excluding hits using strict/automated hit selection criteria.** Sensorgrams for fragments interacting with AChBP with binding levels above the threshold, but not selected as hits due to secondary binding behaviours. These fragments have later been validated as ligands, using X-ray crystallography (FitzGerald and Danielson, manuscript.

#### Supplementary Tables

| Protein | $K_D$ (μM) | |
|---|---|---|
| | **High affinity site** | **Low affinity site** |
| $PTP1B_{1-301}$ | $1.0 \pm 1.1$ | $44 \pm 1.6$ |
| $PTP1B_{1-393}$ | $5.1 \pm 0.2$ | $61 \pm 2.5$ |

**Supplementary Table 1.** $K_D$ values for suramin interactions with $PTP1B_{1-301}$ and $PTP1B_{1-393}$ determined by SPR analysis and assuming a 2:1 stoichiometry with a high and a low affinity site (See Supplementary Note 2 and Supplementary Figure 2.).

| Target protein | Buffer solution |
|---|---|
| tcFPPS, tbFPPS, hFPPS | 10 mM sodium acetate pH 5.0 |
| $PTP1B_{1-301}$, $PTP1B_{1-393}$ | 10 mM sodium acetate pH 5.5 + 1 mM DTT |
| tau K18(C291S, C322S) | 10 mM sodium borate pH 8.5 |
| AChBP | PBS-P+ pH 7.4 |
| LSD1, LSD1/COREST | 10 mM sodium acetate pH 5.0 |

**Supplementary Table 2.** Buffers used for immobilization. DTT was used in buffers for the PTP1B constructs to protect the active-site Cys from oxidation.

| Experiment | NS | SW, ppm | FID | Extra parameters | PULPROG |
|---|---|---|---|---|---|
| 1D $^{1}H$ | 64 | 15.5 | 32768 | - | zgesgp |
| 1D $^{1}H$ CPMG-filtered | 256 | 16 | 32768 | τ = 50 and 800 ms | bdcpmgesdf.rh |
| 2D $^{1}H$-$^{15}N$ SF-HMQC | 32 | F1 = 31.8, F2 = 16 | F1 = 128, F2 = 2048 | NUS = 50% | sfhmqcf3gpph |

**Supplementary Table 3.** Parameters for standard Bruker pulse sequences used to generate 1D and 2D ligand or protein-observed NMR spectra.
